# Human cytomegalovirus protein pUL36: a dual cell death pathway inhibitor

**DOI:** 10.1101/2020.02.03.928564

**Authors:** Alice Fletcher-Etherington, Luis Nobre, Katie Nightingale, Robin Antrobus, Jenna Nichols, Andrew J. Davison, Richard J. Stanton, Michael P. Weekes

## Abstract

Human cytomegalovirus (HCMV) is an important human pathogen and a paradigm of intrinsic, innate and adaptive viral immune evasion. Here, we employed multiplexed tandem mass tag-based proteomics to characterise host proteins targeted for degradation late during HCMV infection. This approach revealed that mixed lineage kinase domain-like protein (MLKL), a key terminal mediator of cellular necroptosis, was rapidly and persistently degraded by the minimally passaged HCMV strain Merlin but not the extensively passaged strain AD169. The strain Merlin viral inhibitor of apoptosis pUL36 was necessary and sufficient both to degrade MLKL and to inhibit necroptosis. Furthermore, mutation of pUL36 Cys^131^ abrogated MLKL degradation and restored necroptosis. As the same residue is also required for pUL36-mediated inhibition of apoptosis by preventing proteolytic activation of pro-caspase 8, we define pUL36 as a multifunctional inhibitor of both apoptotic and necroptotic cell death.

**SIGNIFICANCE STATEMENT:** Cell death is a key defence against viral infection, preventing spread from infected to uninfected cells. Correspondingly, certain viruses encode inhibitors of apoptotic and necroptotic cell death pathways in order to facilitate their persistence. Human cytomegalovirus (HCMV) is an important human pathogen that can block apoptosis, but hitherto it has been unclear whether or how the virus blocks necroptosis. Here, we used a proteomic screen to identify human proteins targeted for destruction by HCMV, finding that the key necroptosis mediator MLKL is degraded throughout infection. MLKL is targeted for degradation by HCMV protein pUL36, which is also instrumental in inhibiting apoptosis. Thus, pUL36 is a dual cell death pathway inhibitor, and may represent an important therapeutic target.

## INTRODUCTION

Human cytomegalovirus (HCMV) infects the majority of the world’s population, with a seroprevalence of 60-80% in Western Europe and the USA, and up to 100% in developing countries (1, 2). Following primary infection, a latent infection is established that persists lifelong. Reactivation from latency to productive infection in immunocompromised people can lead to significant morbidity or mortality, in addition to allograft rejection in transplant recipients (3, 4). Furthermore, HCMV is the commonest infectious cause of congenital disease, affecting ~1/200 pregnancies (5).

Only four drugs are approved by the FDA to treat HCMV, and all are associated with significant toxicity and the threat of drug resistance (2, 6). Small molecule-based disruption of interactions between antiviral host proteins and their viral antagonists could facilitate endogenous inhibition of viral replication (7, 8). Identification and detailed characterisation of such interactions thus has important implications for the development of novel anti-HCMV therapies.

HCMV encodes 170 canonical protein-coding genes (2). A substantial number of additional, non-canonical open reading frames (ORFs) that potentially encode proteins have been identified by ribosomal footprinting and proteomics (9, 10). During productive infection in vitro, expression of HCMV genes is conventionally divided into immediate-early, early and late phases during a ~96 h lytic replication cycle. Recently, five temporal classes of viral protein expression (Tp1-Tp5) have been defined by measuring viral protein profiles over the whole course of infection (11).

Hijacking of the ubiquitin-proteasome system (UPS) to degrade host proteins is ubiquitous across many viral families (12). The proteins degraded by viruses are typically detrimental to viral replication, and can include antiviral restriction factors or components of viral sensing pathways, activating immune cell ligands, and elements of cell death pathways (13, 14). We have shown previously that >900 host proteins are downregulated >3-fold over the course of HCMV infection, with 133 proteins degraded during the early phase, of which 89% are targeted to the proteasome. These data led directly to the identification of candidate natural killer (NK) cell ligands and identified helicase-like transcription factor (HLTF) as a novel antiviral restriction factor (10, 11). However, it is not yet known which proteins are degraded later during infection, or throughout a whole infection time-course. Furthermore, it is unclear whether or how HCMV inhibits necroptotic cell death pathways, which represent a key defence against viral spread within an infected host (15).

Here, a systematic examination of protein degradation at 48 h of HCMV infection determined that degradation of MLKL, a pseudokinase that acts as the non-enzymatic effector of necroptosis (16), was sustained throughout early and late infection. Receptor-interacting serine/threonine-protein kinase 3 (RIP3)-dependent phosphorylation of MLKL drives the transition of monomeric MLKL to a necroptotic oligomer that translocates to and then disrupts the plasma membrane (17, 18). The data showed that MLKL was degraded by the minimally-passaged Merlin strain of HCMV, but not by the highly passaged laboratory-adapted strain AD169. The strain Merlin UL36-encoded viral inhibitor of caspase-8 activation (vICA/pUL36), which is known to function as a potent inhibitor of apoptosis, interacted with MLKL and was necessary and sufficient both to degrade MLKL and to inhibit TNFα-stimulated necroptosis. Whereas HCMV strain AD169 sensitised cells to necroptosis, strain Merlin prevented this sensitisation. pUL36 is thus a multifunctional cell death inhibitor capable of inhibiting both apoptotic and necroptotic pathways.

## RESULTS

### Host proteins targeted for degradation at 48 h of HCMV infection

We applied the proteasomal inhibitor MG132 to build a global picture of proteins degraded late during HCMV infection. TERT-immortalised primary human fetal foreskin fibroblasts (HFFF-TERTs) were infected with HCMV strain Merlin at a multiplicity of infection (MOI) of 10 for 48 h, with application of MG132 (or the equivalent amount of DMSO as a control) for the final 12 h of infection **(Figure 1A)**. MG132 is known to inhibit calpains and lysosomal cathepsins in addition to the proteasome (10), and was used to generate a comprehensive list of proteins targeted for degradation by HCMV rather than to identify proteins specifically degraded by the proteasome. 8,476 human and 186 viral proteins were quantified in two biological replicates (**Dataset S1**, which includes an interactive ‘Plotter’). Two ratios and associated significance values were calculated for each protein: (i) virus:mock infection and (ii) virus+MG132:virus infection **(Figure 1B)**. 52 proteins met high confidence criteria for degradation, with a fold downregulation and rescue of >2 and p<0.05 for both ratios.

**Figure 1.**
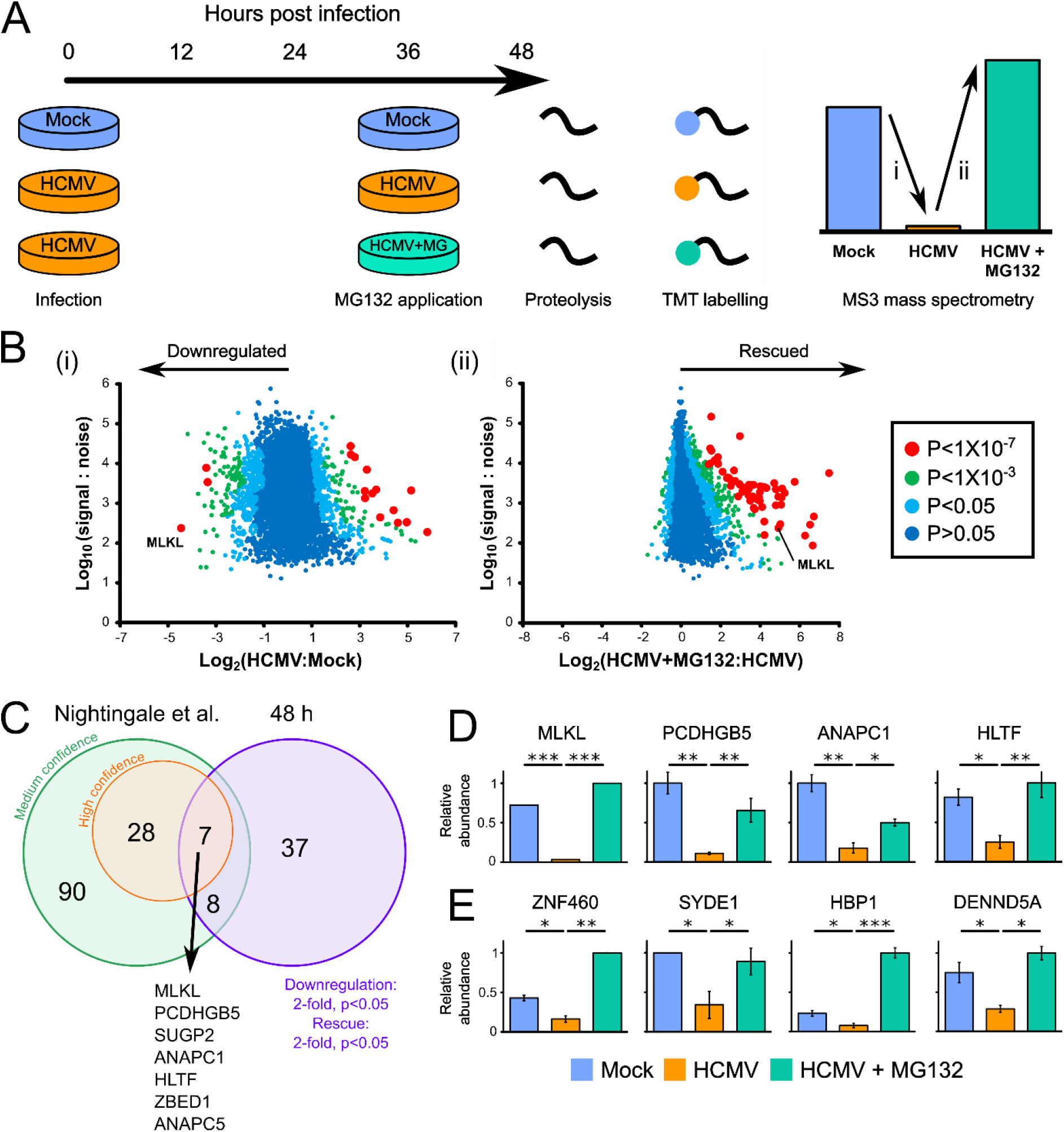
Identification of proteins targeted for degradation by HCMV at 48 hpi. **(A)** Schematic of the experimental method, conducted in biological duplicate. Cellular lysates from the second 48 h biological replicate were analysed simultaneously with residual lysates from the 12 h degradation screen in the Nightingale *et al.* study (10), facilitating a direct comparison (**Figure S1A**). Peptides from each sample were labelled with tandem mass tags and analysed by MS3 mass spectrometry. **(B)** Scatterplots of human proteins quantified at 48 hpi in one or both replicates, showing averaged ratios. P-values were estimated using Significance B values, then corrected for multiple hypothesis testing (20). K-means clustering suggested there were at least nine different patterns of protein expression across the samples **(Figure S1B, Dataset S1C)**. **(C)** Overlap between early degradation data from the Nightingale *et al.* study (using either stringent or sensitive statistical criteria) (10) and 48 h degradation data presented in this study. **(D)** Examples of the seven proteins degraded with high confidence both early and late during infection. MLKL was only quantified in one replicate. Error bars: range. P-values were calculated as described in (B). *p<0.05, **p<0.001, ***p<1×10^−7^. Additional proteins were likely to have been degraded both early and late during infection **(Dataset S1, Figure S1A)**, but did not pass the stringent filtering criteria used at one or other of the time points. **(E)** Examples of the 37 proteins degraded at 48 h that were not confidently degraded at early time points. These comprised: (i) proteins not degraded at early time points (shown in this figure), (ii) proteins that were insufficiently degraded at early time points to pass filtering criteria, and (iii) proteins not quantified in the Nightingale *et al.* study (10). Additional examples and the corresponding 12 h data are shown in **Figure S1A**.

From a comparison with our previous study of HCMV-induced protein degradation between 2-24 h (10), seven proteins were degraded with high confidence throughout early and late infection **(Figures 1C, 1D)**. These proteins included HLTF and Anaphase Promoting Complex subunits 1 and 5 (ANAPC1/5), whose degradation by HCMV has been well characterised (10, 19). The effector of necroptosis MLKL was the most significantly downregulated protein at 48 h of HCMV infection and was among proteins most significantly rescued by addition of MG132 **(Figure 1B)**. In comparison, 37 proteins degraded at 48 h did not score highly for degradation at early time points (10) **(Figures 1C, 1E, S1A)**.

### Degradation of MLKL is mediated by immediate early protein pUL36

We previously took a systematic approach to identifying the viral proteins responsible for the degradation of host factors, employing a panel of recombinant viruses deleted for various blocks of HCMV genes that are non-essential for replication *in vitro* (10). These viruses included strain AD169, a highly passaged, laboratory-adapted strain that contains a deletion in the U_L_/*b’* region (encoding 20 canonical genes, UL133-UL150A), frameshifts in RL5A, RL13 and UL131A, and a non-synonymous substitution in UL36 that inactivates the ability of pUL36 to bind procaspase 8 and inhibit apoptosis (21, 22). MLKL was downregulated by strain Merlin viruses but not by strain AD169, and this was confirmed by immunoblotting **(Figures 2A-B)**. This indicated that one or more differences in AD169 abrogate virus-mediated MLKL degradation. To identify viral factors interacting with MLKL, a SILAC immunoprecipitation (IP) was performed in HCMV strain Merlin-infected HFFF-TERTs stably expressing MLKL tagged with HA at the C-terminus. pUL36 co-precipitated with MLKL, and this was confirmed by a reciprocal IP **(Figures 2C-D, Dataset S2)**. This interaction was further supported by an immunofluorescence study showing cytoplasmic co-localisation between MLKL-HA and pUL36 in stably expressing HFFF-TERTs **(Figure S2A)**. To determine whether pUL36 interacts with inactive, unphosphorylated MLKL or active, phosphorylated MLKL we re-searched the pUL36-V5 SILAC IP data (**Figure 2D**) using a variable phospho-modification. All identified MLKL peptides were unphosphorylated, and encompassed all known sites of activating phospho-modifications **(Figure S3)**. This suggests that pUL36 interacts with unphosphorylated MLKL but does not exclude an additional interaction with phosphorylated MLKL.

**Figure 2.**
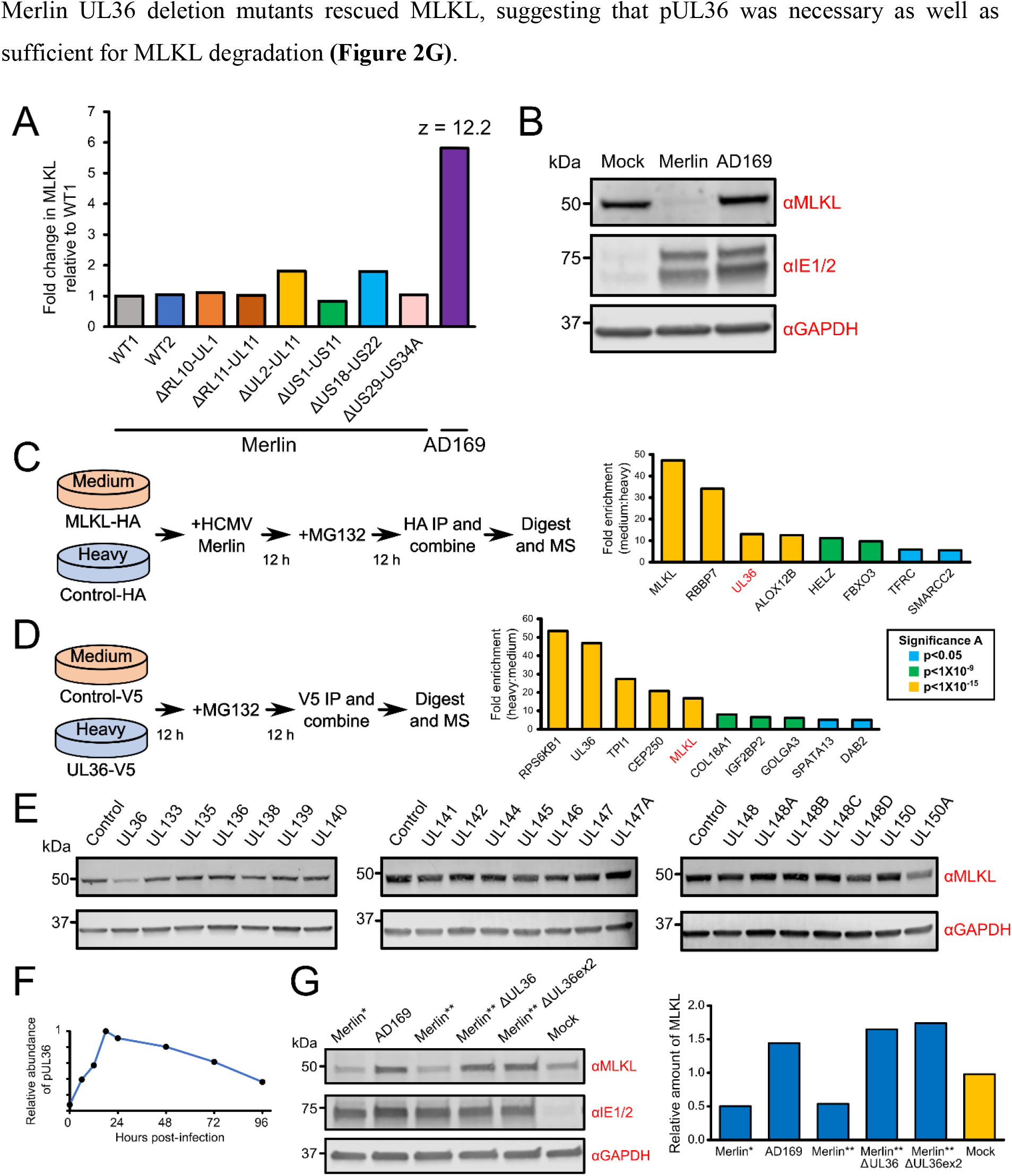
HCMV strain Merlin pUL36 is necessary and sufficient to degrade MLKL. **(A)** Relative abundance of MLKL from a proteomic screen of >250 proteins published in the Nightingale *et al.* study (10). HFFF-TERTs were infected with strain Merlin (WT1), WT1 that lacked UL16 and UL18 (WT2), one of six block-deletion viruses derived from WT1 or WT2, or strain AD169 (MOI = 10, 72 h infection). A z-score of >5 was considered significant. **(B)** Immunoblot confirming that HLTF is downregulated by strain Merlin but not by strain AD169 (MOI = 5, 48 h infection, cells lysed in RIPA buffer). MLKL was similarly downregulated in cells lysed in 2% SDS (**Figure S2E**). This was consistent with the observed MLKL downregulation in cells lysed with 6 M guanidine for proteomic experiments (**Figure 1**), and indicated that MLKL was degraded as opposed to being translocated to RIPA-insoluble membrane-associated complexes. **(C)** SILAC immunoprecipitation of C-terminally HA-tagged MLKL or control in the presence of strain Merlin infection (MOI = 3, 24 h infection in the presence of 10 μM MG132 for the final 12 h). Proteins enriched >5-fold are shown. P-values were estimated using the method of significance A and corrected for multiple hypothesis testing (20). **(D)** SILAC immunoprecipitation of C-terminally V5-tagged pUL36 or control in the presence of 10 μM MG132 for 12 h. Proteins that were enriched >5-fold are shown, and p-values were estimated as described in (C). **(E)** Immunoblot showing that pUL36 is sufficient to downregulate MLKL. A series of HFFF-TERT cell lines stably expressing genes in the UL133-150A region were lysed in RIPA buffer and analysed by immunoblotting. Confirmation of viral protein expression was achieved by immunoblotting or mass spectrometry (23) (**Dataset S3D, Figure S2B)**, except for pUL136, which was not detected by either method. MLKL was similarly downregulated by pUL36 in cells lysed in 2% SDS (**Figure S2F**, as shown during HCMV infection in **Figure S2E)**. **(F)** Temporal profile of strain Merlin pUL36 in whole-cell lysates harvested at different times over the whole course of infection, from the Weekes *et al.* study (11). **(G)** Immunoblot showing that pUL36 is necessary for downregulation of MLKL (MOI = 5, 24 h infection). Cells were infected with WT strain Merlin (Merlin*), a version of Merlin in which intact genes UL128 and RL13 are under tet regulation (Merlin**), two UL36 deletion viruses derived from Merlin**, or strain AD169. ∆UL36ex2 has a deletion in exon 2. Right panel: the relative amount of MLKL normalised to GAPDH. Overall this data suggests that pUL36 is necessary and sufficient to degrade MLKL.

A series of HFFF-TERT cell lines stably expressing strain Merlin UL36 or each of the individual genes in the UL133-UL150A region were next screened to determine whether any other viral protein contributed to MLKL degradation. Expression of UL36 was sufficient to reduce the level of MLKL by 3.7-fold **(Figures 2E, S2B-D)**. MLKL expression was not modulated more than 2-fold by any other proteins in the U_L_/*b’* region. Despite substantial variation in the level of expression of some of the viral proteins, this did not correlate with relative MLKL abundance (**Figure S2D**). pUL36 could be detected from 6 hpi, and was expressed with Tp2 (temporal protein profile 2) kinetics, matching the kinetics of MLKL degradation (11) **(Figure 2F)**. Finally, infection with strain AD169 or two independent strain Merlin UL36 deletion mutants rescued MLKL, suggesting that pUL36 was necessary as well as sufficient for MLKL degradation **(Figure 2G)**.

### pUL36 protects cells from necroptosis

Activation of death receptors such as TRAILR1/2, TNFR1 and Fas leads to apoptotic cell death via an activating cleavage of pro-caspase-8 (24). In the presence of caspase inhibition or limiting levels of ATP, extrinsic apoptosis shifts towards the necroptotic pathway. This is dependent on an interaction between Receptor Interacting Serine/Threonine Kinases 1 and 3 (RIP1/3) through their homotypic interaction motif (RHIM) domains (25) (**Figure 3A**). Necroptosis can also be activated by cytoplasmic sensing of murine cytomegalovirus (MCMV) DNA by Z-DNA-binding protein 1 (ZBP1), or ligation of toll-like receptors 3 or 4 (TLR3/4), which activate RIP3 through alternative RHIM domain-containing adaptors (25). All three pathways converge with RIP3-dependent phosphorylation and activation of MLKL, which transitions from an inactive monomer to a necroptotic oligomer. The oligomer translocates to and disrupts the plasma membrane, leading to cell swelling and loss of plasma membrane integrity, although the exact mechanism of membrane disruption is unclear (17, 18, 26) (**Figure 3A**).

**Figure 3.**
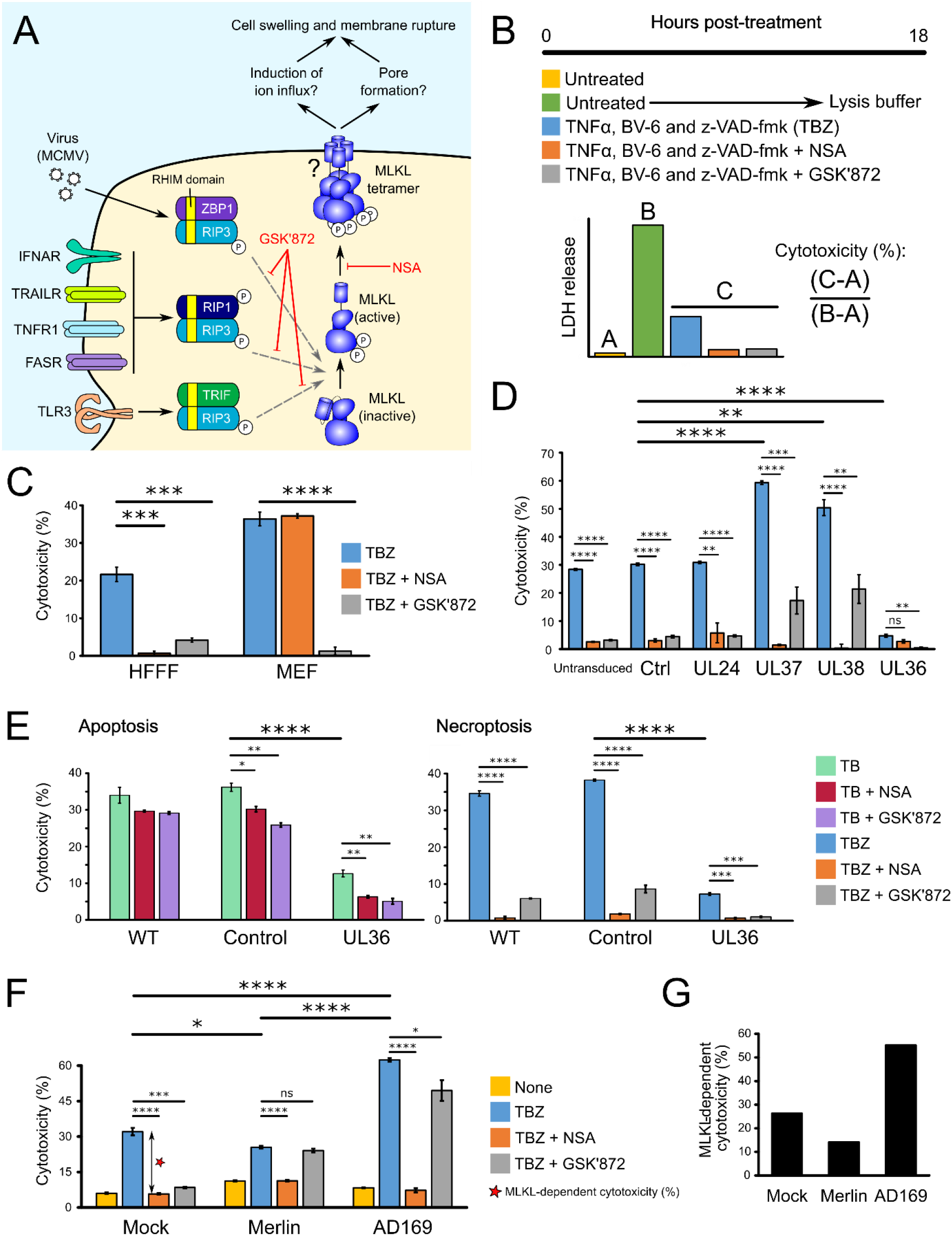
Role of MLKL degradation in HCMV infection. **(A)** The necroptosis pathway and the mechanisms of action of small molecule necroptosis inhibitors. **(B)** Schematic of the necroptosis assay, which was performed in biological triplicate conducted in parallel and repeated in 2 or 3 completely independent experiments. Cytotoxicity was quantified by measuring lactate dehydrogenase (LDH) release using the CytoTox 96® Non-Radioactive Cytotoxicity Assay (Promega). Cytotoxicity was calculated as a percentage of 100% lysis (from untreated cells lysed with lysis buffer) after subtraction of background LDH release from live cells. To stimulate apoptosis, cells are treated with TB instead of TBZ. **(C)** TBZ induces MLKL- and RIP3-dependent necroptosis, which was inhibitable by GSK’872 in HFFF-TERTs and MEFs. NSA inhibited necroptosis in HFFF-TERTs but not MEFs, likely because NSA does not inhibit murine forms of MLKL (17). Error bars show standard error of the mean (SEM). P-values were estimated using a two-tailed t-test (n=3). ***p < 0.001, ****p < 0.0001. Data are representative of three independent experiments. To dissect fully the conditions of this assay, stimulation with each of T, B and Z alone or in combination was examined in the presence or absence of inhibitors (**Figure S4B-C**). HFFFs were insensitive to T or Z alone, TZ or BZ. Cell death was stimulated by B alone in the absence of death receptor stimulation, an effect that has been observed in other cell types (40) **(Figure S4B)**. The cell death initiated by BV6 was MLKL- and RIP3-independent, but also caspase-8-dependent (inhibited by the addition of Z), suggesting that BV6 alone was not responsible for the necroptotic cell death observed in the TBZ condition **(Figures S4B-C)**. Collectively, these results suggest that only in the presence of TBZ was cell death dependent on MLKL and RIP3. **(D)** pUL36 is sufficient to inhibit necroptosis in stably transduced HFFF-TERTs treated with TBZ ± inhibitors. In addition to untransduced cells, two control vectors were employed. ‘Ctrl’ cells were transduced with a vector containing a short, randomised DNA sequence. pUL24 was included as a control HCMV tegument protein that has a similar size to pUL36 but lacks any known role in cell death. Error bars: SEM. P-values were estimated using a two-tailed t-test (n=3). **p < 0.01, ***p < 0.001, ****p < 0.0001. Data are representative of two independent experiments. **(E)** pUL36 inhibits both apoptosis and necroptosis. WT, control or pUL36-expressing HFFF-TERTs were treated for 18 h with either TB or TBZ to stimulate apoptosis or necroptosis, respectively, in the presence or absence of NSA or GSK’872. Error bars: SEM. P-values were estimated using a two-tailed t-test (n=3). *p < 0.05, **p < 0.01, ***p < 0.001, ****p < 0.0001. Data are representative of two independent experiments. **(F)** Functional pUL36 was required for inhibition of necroptosis, comparing 48 h mock, Merlin or AD169 infection (MOI = 5) in HFFF-TERTs subsequently stimulated with TBZ ± inhibitors for 18 h. Baseline cytotoxicity in untreated cells was not subtracted from the other values and is shown as a separate yellow bar. Error bars: SEM. P-values were estimated using a two-tailed t-test (n=3). *p < 0.05, ***p < 0.001, ****p < 0.0001. Data are representative of two independent experiments. **(G)** MLKL-dependent cytotoxicity was calculated from the difference between % cytotoxicity (TBZ alone) versus (TBZ+NSA), shown by the double-headed arrow in **Figure 3F**.

The HCMV UL36-encoded viral inhibitor of caspase-8 activation (vICA/pUL36) inhibits apoptosis by binding the pro-domain of caspase-8 and preventing its proteolytic activation (22). We sought to determine whether Merlin pUL36 additionally inhibits necroptosis. *In vitro*, necroptosis can be stimulated by a combination of TNFα (T), BV6 (B, an IAP antagonist that sensitises cells to TNFα-induced cell death), and the pan-caspase inhibitor Z-VAD-fmk (Z) **(Figure 3B)**(27, 28). Although previous reports have suggested that HFFFs are not susceptible to necroptosis due to limiting levels of RIP3 (29), both RIP3 and RIP1 were detectable in HFFF-TERTs by proteomics (**Figure S4A**). Necroptosis was induced by T+B+Z (TBZ) in HFFF-TERTs as well as in immortalised mouse embryonic fibroblasts (MEFs), which are highly susceptible to necroptosis (30) and were therefore used as a positive control **(Figure 3C)**. Two inhibitors were employed to determine whether the stimulated death pathway was canonical MLKL- and RIP3-dependent necroptosis. GSK’872 binds to and inhibits the RIP3 kinase domain (31, 32), whereas necrosulfonamide (NSA) inhibits downstream effector functions of MLKL via covalent reaction with the Cys86 residue of human but *not* murine MLKL (17, 33). NSA and GSK’872 inhibited TBZ-induced cytotoxicity in HFFF-TERTs, indicating that these cells expressed sufficient RIP3 to induce measurable canonical necroptosis **(Figure 3C)**.

Omoto *et al*. previously demonstrated that pUL36 does not impact the ability of HCMV strain Towne to protect against necroptosis in fibroblasts stably transduced with RIP3 (29). In contrast to this conclusion, we found that strain Merlin pUL36 was sufficient to inhibit necroptosis. The two other HCMV-encoded inhibitors of apoptosis, pUL37×1/vMIA (viral mitochondria-localized inhibitor of apoptosis) and pUL38 (34, 35), augmented MLKL-dependent necroptosis (as demonstrated by complete inhibition by NSA). However, cell death was incompletely inhibited by GSK’872, suggesting that a RIP3-independent mechanism might be acting in addition **(Figure 3D)**. Next, WT, control and UL36-expressing cell lines were treated with TB or TBZ to assess the effect of pUL36 on apoptosis and necroptosis respectively **(Figure 3E)**. Other inhibitors of caspase-8, including MCMV pM36 and the compound z-VAD-fmk (Z), do not inhibit death receptor-stimulated cell death and instead activate necroptosis (36–39). In contrast, pUL36 was able to inhibit cell death stimulated by TB (in agreement with Skaletskaya *et al.* (22)), however the residual cell death was partly RIP3 and MLKL-dependent **(Figure 3E)**. These data suggest that, in the presence of TB, pUL36 partly converts apoptotic cell death to necroptosis, and inhibits both apoptosis and necroptosis. In the presence of TBZ, apoptosis is potently inhibited by the addition of Z, and pUL36 inhibits purely necroptotic cell death.

Finally, infection of HFFFs with HCMV strains Merlin and AD169 for 48 h prior to TBZ stimulation suggested that inhibition of MLKL-dependent necroptosis required functional pUL36. Pre-infection with Merlin had a slight protective effect on necroptosis, whereas AD169 infection amplified the effect of TBZ **(Figures 3F-G)**. Cell death induced by TBZ after infection with HCMV also appeared to have a RIP3-independent component, as GSK’872 had little inhibitory effect.

### Substitution of Merlin pUL36 Cys131 abrogates inhibition of necroptosis by abolishing MLKL binding and degradation

Strain Merlin and strain AD169 pUL36 differ by five amino acid residues **(Figure S5)**, including a Merlin Cys^131^ **→** AD169 Arg^131^ substitution. This single replacement abrogates inhibition of apoptosis, as pUL36 can no longer bind procaspase-8 (22). Five Merlin pUL36 mutants corresponding to the five amino acid substitutions between strain Merlin and strain AD169 were constructed in order to determine which were important for degradation of MLKL. Only the C131R substitution prevented pUL36 from binding and downregulating MLKL (**Figure 4A-B**). The same pUL36 mutant was also unable to protect cells from necroptosis, suggesting that this single residue plays a key functional role in inhibition of both apoptosis and necroptosis by pUL36 **(Figure 4C)**.

**Figure 4.**
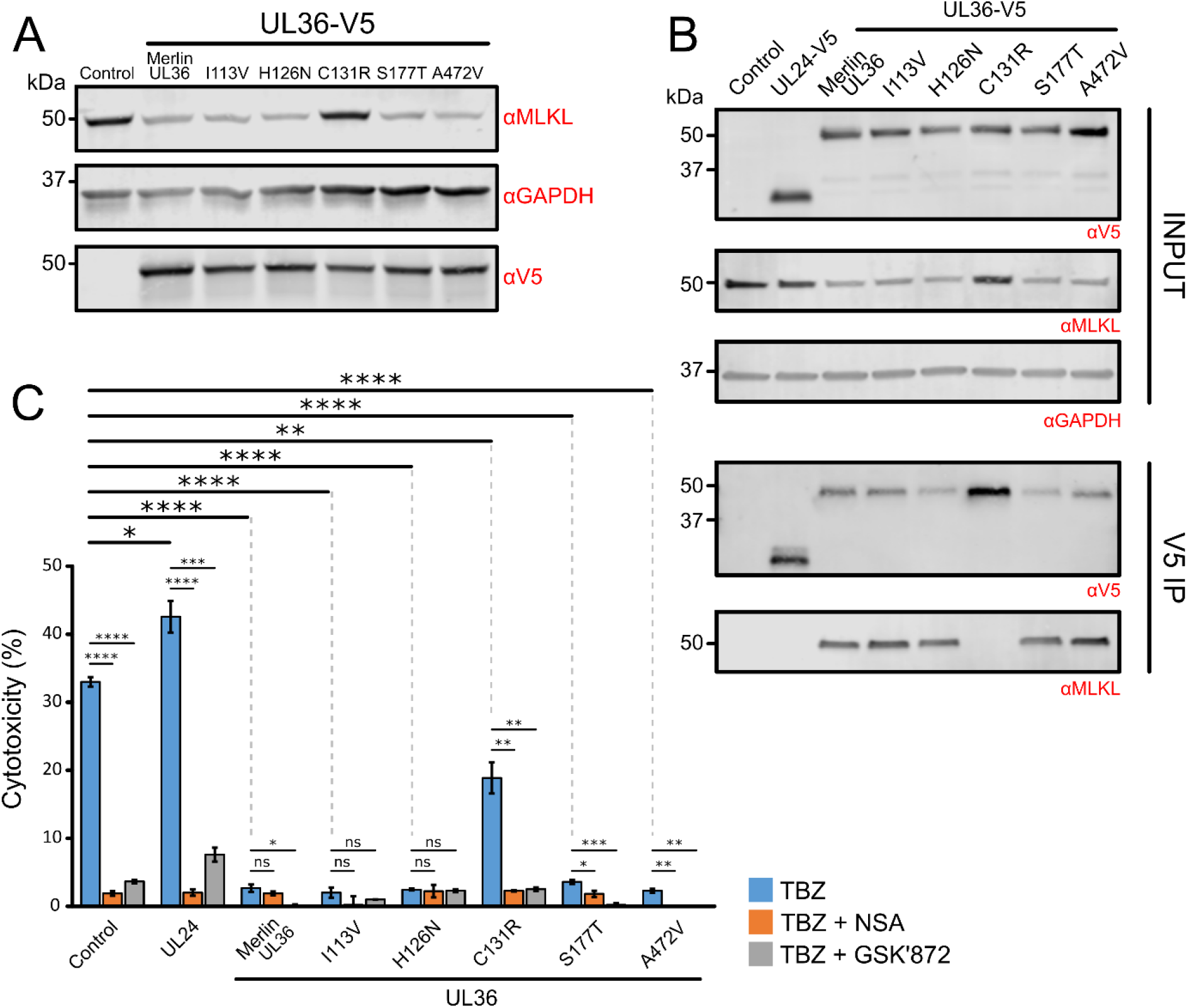
Substitution of Cys^131^ in strain Merlin pUL36 abrogates binding to and degradation of MLKL, and inhibition of necroptosis. **(A)**Immunoblot showing that downregulation of MLKL is dependent on the pUL36 Cys^131^ residue. HFFF-TERTs were stably transduced with the indicated C-terminally V5-tagged UL36 constructs. Individual point mutations corresponding to the five amino acid substitutions between strain Merlin and strain AD169 pUL36 are shown above the blot and detailed in **Figure S5**. **(B)** Interaction between pUL36 and MLKL is dependent on pUL36 Cys^131^, shown by a V5 co-IP in the cell lines described in (A). Immunoblots were probed with anti-V5, anti-MLKL and anti-GAPDH antibodies. **(C)** Percentage cytotoxicity of cell lines described in (A) that were treated with TBZ ± NSA or GSK’872. Error bars: SEM. P-values were estimated using a two-tailed t-test (n=3). *p<0.05, **p < 0.01, ***p < 0.001, ****p< 0.0001. Data are representative of two independent experiments. Low-level variation in sensitisation to necroptosis was observed between the unmodified, control and UL24-expressing cell lines (comparing data in this figure to **Figure 3D**).

## DISCUSSION

Herpesviruses persist lifelong in infected individuals by comprehensive modulation of adaptive and innate immunity. Multiple viral proteins are deployed to target host factors for degradation, many very early during infection (10). The present study provides a systematic, searchable database that examines host protein degradation at 48 h of HCMV strain Merlin infection, approximately 50-66% of the way through the lytic replication cycle in HFFFs. In addition to identifying proteins degraded throughout the HCMV replication cycle, these data may be particularly useful for identifying which viral protein targets a given host factor for degradation. We defined previously the kinetics of expression of the majority of canonical HCMV proteins, which can now be compared to the kinetics of host protein degradation, from 6 to 48 h post-infection (10, 11). Other mechanisms of host protein degradation that are not inhibited by MG132 are also likely to be subverted by HCMV. However, 87-89% of degraded proteins were targeted to the proteasome by 24 h post-infection (10).

The key roles of necroptosis in protecting cell populations against virus infection are highlighted by the impressive range of known viral countermeasures. Necroptosis inhibitors are particularly widespread amongst herpesviruses, with MCMV and herpes simplex viruses (HSV) types 1 and 2 encoding conserved RHIM-domain containing proteins (M45, ICP6 and ICP10, respectively) which compete with host RHIM-domain adaptor proteins such as RIP1 for binding to RIP3 (41–43). The HCMV M45 ortholog pUL45 does not contain a RHIM-domain and does not inhibit cell death (44). Instead, it had been suggested that HCMV targets the necroptotic pathway downstream of RHIM signalling, after RIP3-dependent phosphorylation of MLKL, although the mechanism had not been elucidated (29).

As other viral inhibitors of caspase-8 activation have been shown to sensitise cells to programmed necrosis (39, 45), it has been assumed that HCMV pUL36 would have a similar effect, requiring HCMV to encode a separate mechanism of necroptosis inhibition in order to evade cell death completely. However, although MCMV M36 can sensitise cells to necroptosis (39), the same phenomenon has not been demonstrated for HCMV pUL36, which is not surprising given that both proteins exhibit only 19% sequence identity (46). Furthermore, the question of why HCMV pUL36 expression can inhibit death receptor-stimulated apoptosis instead of stimulating necroptosis as per the action of other inhibitors of caspase-8 such as z-VAD-fmk and MCMV M36 (36–39) had not hitherto been examined. Here, we explain these observations by showing that HCMV pUL36 mediates degradation of MLKL.

pUL36 can bind both pro-caspase-8 (22) and MLKL, facilitating potent inhibition of both apoptosis and necroptosis. Indeed, these two functions can be inhibited by the same single amino acid substitution gained during passage of HCMV in cell culture (21, 22). This suggests that a larger cell death complex may form from the caspase-8:FADD:RIP1 FADDosome/Ripoptosome (47–49) and the RIP1:RIP3 necrosome (25), enabling an interaction between pUL36, caspase-8 and MLKL.

Necroptotic activation of MLKL can influence many other cellular processes, including inflammasome activation, endosomal trafficking, extracellular vesicle generation and autophagy (50–53). It is therefore possible that HCMV-mediated degradation of MLKL may have other consequences for viral pathogenesis, including effects on virion assembly, trafficking and cell-to-cell spread.

Previous studies investigating the interaction between HCMV and necroptosis have employed HFFFs stably transduced with RIP3 in order to confer susceptibility to necroptosis, which can otherwise be lost during cell propagation (29). Using these cells, the authors found that all tested HCMV strains, including AD169, were able to inhibit TNFα-stimulated necroptosis, although strain Merlin inhibited necroptosis more potently than AD169, consistent with our results **(Figure 3F)**. Furthermore, comparison of a WT strain Towne (encoding pUL36 with a cysteine at position 131) with a Towne ∆UL36 mutant suggested that pUL36 was necessary for inhibition of apoptosis but *not* necroptosis (29). In addition to mediating cell death, RIP3 has been implicated in NF-kB and inflammasome activation and can induce apoptosis when overexpressed (32, 54). It is possible that overexpression of RIP3 has off-target effects, which may explain the discrepancy between the previously published work and the present study; the experimental setting may clearly influence the outcome observed with pUL36. In addition, while the Omoto *et al*. study suggested that HCMV targets necroptosis downstream of MLKL phosphorylation, all MLKL peptides in our pUL36-V5 SILAC IP were unphosphorylated **(Figures 2D and S3)**, suggesting that pUL36 affects the monomeric MLKL pool. However, this does not exclude the potential for additional HCMV-mediated direct or indirect mechanisms of necroptosis inhibition. Further evidence that results can be dependent on cell type and the presence of RIP3 overexpression came from use of the Towne ∆UL36 mutant to show that pUL36 can inhibit caspase-independent cell death during late stages of macrophage differentiation (55). This also suggests that pUL36-mediated degradation of MLKL may occur in more than one cell type. The HFFF-TERT cell line used in the present study is susceptible to RIP3 and MLKL-dependent canonical necroptosis **(Figure 3C)** and may be an invaluable resource for future studies of viral modulation of cell death.

Infection of HFFF-TERTs with HCMV prior to TBZ stimulation resulted in induction of a form of cell death that was not completely inhibited by GSK’872, which is suggestive of an RIP3-independent but MLKL-dependent mechanism **(Figure 3F)**. A similar but less significant effect was observed in cells expressing the HCMV apoptosis inhibitors pUL37 and pUL38 **(Figure 3D)**. RIP3-independent necroptosis in fibroblasts has been reported previously by others (30) but remains poorly characterised. Infection with HCMV strain AD169, which lacks a functional pUL36 protein, sensitised cells to necroptosis **(Figure 3F)**, in accord with the observed increase in MLKL protein upon infection with viruses deficient in functional pUL36 **(Figure 2G)**. This is likely to be due to IFN-mediated upregulation of MLKL (56). Strain Merlin protein pUL36 was able to counteract the effect of necroptosis sensitisation, rather than abrogating necroptosis entirely **(Figure 3F)**.

HCMV pUL36 and its orthologs belong to the US22 family of herpesvirus proteins, which characteristically feature a set of motifs I to IV (46). Cys^131^, which is found within motif III, is conserved within the primate cytomegaloviruses but is replaced by conservative substitutions in rodent cytomegalovirus orthologs that do not abrogate apoptosis inhibition. In addition, the MCMV ortholog pM36, which displays only 19% sequence similarity to HCMV pUL36 and lacks the region that includes Cys^131^, is still an efficient suppressor of apoptosis (46). This suggests that the Arg^131^ residue of strain AD169 pUL36 may restrict cell death inhibition, as opposed to Cys^131^ being required for this function.

Only four drugs are currently available for treating HCMV, all exhibiting significant adverse effects and the potential of drug resistance. The identification of a potentially inhibitable interaction between a single residue of a viral antagonist and key mediators of both necroptosis and apoptosis may therefore be of substantial therapeutic significance. In addition to pUL36-MLKL, other interactions involving distinct antiviral pathways could be targeted simultaneously to potently inhibit viral replication, for example between HCMV pUL145 and the recently identified restriction factor HLTF (10). Moreover, our data are likely to identify further proteins that have roles in restricting infection by HCMV or other viruses.

## MATERIALS AND METHODS

Extended materials and methods can be found in the supplementary information (SI)

### Viral infections

The required volume of viral stock to achieve the multiplicity of infection (MOI) described in the results section was diluted in serum-free Dulbecco’s modified Eagle’s medium (DMEM), mixed gently and applied to HFFF-TERTs. Mock infections were performed identically but with DMEM instead of viral stock. Time zero was considered the time at which cells first came into contact with virus. Cells were incubated with virus for 2 h at 37 °C on a rocking platform, and then the medium was replaced with DMEM + 10% FBS.

### Proteomic Screen

HFFFs-TERTs were infected as described above in biological duplicate at an MOI of 10. At 36 hpi, 10 μM MG132 or the equivalent volume of DMSO was added to the cells for the last 12 h of infection. Samples from the second replicate were digested and analysed with residual samples from the 12 h degradation screen from the Nightingale *el al.* study (10). Methods for whole cell lysate protein preparation and digestion, peptide labelling with tandem mass tags, HpRP fractionation, liquid chromatography-mass spectrometry and data analysis are discussed in detail in the Nightingale *et al.* study (10) and recapitulated in the SI.

### Immunoblotting

Cells were lysed, sonicated and clarified by centrifugation, and protein concentration was measured using a bicinchoninic acid (BCA) assay. Samples were denatured and reduced, and then the proteins were separated by SDS polyacrylamide gel electrophoresis (PAGE), transferred to a polyvinylidene difluoride (PVDF) membrane (0.45 μm pore), and probed using the primary and secondary antibodies detailed in the SI. Fluorescent signals were detected using the Odyssey CLx Imaging System (LI-COR), and images were processed and quantified using Image Studio Lite V5.2 (LI-COR).

### Plasmid construction and transduction

Lentiviral expression vectors encoding MLKL-HA, the V5-tagged viral proteins pUL36 and pUL133-pUL150A and controls, were synthesised by PCR amplification of the genes and cloning them into Gateway vectors (57). V5-tagged UL36 point mutants were generated by PCR site-directed mutagenesis. The primers and templates used are described in the SI. Stable cell lines were generated by transduction with lentiviruses produced via the transfection of HEK293T cells with lentiviral expression vectors and helper plasmids.

### Immunoprecipitation

Cells were harvested in lysis buffer, tumbled on a rotator and then clarified by centrifugation and filtration. After incubation with immobilised mouse monoclonal anti-V5 or anti-HA agarose resin, samples were washed and then subjected either to immunoblotting or mass spectrometry (see SI).

### Cell death assays

For assays in the absence of infection, 96-well plates were seeded with HFFF-TERTs or immortalised MEFs and incubated for 24 h. Cells were incubated for 18 h with 30 ng/ml TNFα, 5 μM BV-6 and/or 25 μM z-VAD-fmk in the presence or absence of 0.5 μM necrosulfonamide (NSA) or 1.5 μM GSK’872, or DMSO alone (control). Half of the control cells were lysed to measure lactate dehydrogenase (LDH) release from maximally lysed cells. The other half were used to measure background LDH release from live, untreated cells. LDH release was measured using the CytoTox 96® Non-Radioactive Cytotoxicity Assay. Average absorbance values derived from culture medium alone were subtracted from each absorbance value from experimental wells. For assays in the presence of infection, HFFF-TERTs were seeded into 96-well plates, incubated for 24 h, and infected with HCMV (MOI = 5). The medium was changed 48 h after infection to stimulate necroptosis as described above.

## ACKNOWLEDGEMENTS

We are grateful to Prof. Steve Gygi for providing access to the “MassPike” software pipeline for quantitative proteomics. This work was supported by a Wellcome Trust Senior Clinical Research Fellowship (108070/Z/15/Z) to MPW, an MRC Project Grant (MR/P001602/1) to RJS, and an MRC Programme Grant (MC_UU_12014/3) to AJD. This study was additionally supported by the Cambridge Biomedical Research Centre, UK.

