## Supplemental Text and Figures for "Human cytomegalovirus protein pUL36: a dual cell death pathway inhibitor"

**This PDF file includes:**

Supplementary text (Materials and Methods)
Figures S1 to S5
Legends for Datasets S1 to S3
SI References

**Other supplementary materials for this manuscript include the following:**

Datasets S1 to S3

**Supplementary Information Text**

### MATERIALS AND METHODS

#### Cells and cell culture

All fibroblast cell lines generated in this paper were derived from human foetal foreskin fibroblasts immortalised with human telomerase (HFFF-TERTs) (1). HFFF-TERTs have been tested at regular intervals since isolation to confirm that human leukocyte antigen (HLA) and MHC Class I Polypeptide-Related Sequence A (MICA) genotypes, cell morphology and antibiotic resistance are unchanged. HFFF-Tets (used for growing tet-regulated viruses) additionally constitutively expressed the tetracycline (tet) repressor. Mouse embryonic fibroblasts (MEFs) immortalised with SV40 large T antigen were kindly provided by Dr Evan Reid (University of Cambridge) and are described in the Allison *et al.* study (2). HEK293T cells used for generation of lentivirus were obtained as a gift from Professor Paul Lehner (University of Cambridge). All cells were grown at 37 °C in 5 % v/v $CO_2$ in Dulbecco's modified Eagle's medium (DMEM) supplemented with foetal bovine serum (FBS: 10% v/v), and 100 IU/ml penicillin / 0.1 mg/ml streptomycin (DMEM/FBS/PS). For SILAC immunoprecipitations (IPs), cells were grown for seven passages in SILAC DMEM (Gibco), which was supplemented with 10% dialysed FBS (Gibco), penicillin/streptomycin, 280 mg/l L-proline (Sigma-Aldrich), and either medium (Arg 6, Lys 4) or heavy (Arg 10, Lys 8) amino acids (CK Isotopes) at 50 mg/ml.

#### Viruses

The genome sequence of HCMV strain Merlin (GenBank accession AY446894) is designated the reference HCMV sequence (RefSeq accession NC_006273.2) by the National Center for Biotechnology Information (3, 4). A recombinant version (RCMV1111) of this strain was derived by transfection of a sequenced BAC clone (4). RCMV1111 contains point mutations in two genes (RL13 and UL128) that enhance replication in fibroblasts (4). The block HCMV deletion mutants are described in the Nightingale *et al.* study (5). The AD169-GFP virus (RCMV288) is described in the McSharry *et al.* study (1). RCMV1502 is an RCMV1111 recombinant that has tet-operators 5' to the RL13 and UL128 coding sequences (Merlin-$RL13_{tetO}$-$UL128_{tetO,}$) and is described in the Stanton *et al.* study (4). UL36 deletion mutants were generated by recombineering RCMV1502 (4). Two deletion mutants were generated, one lacking the whole UL36 coding sequence (ΔUL36; RCMV2288), and the other lacking the second UL36 exon (ΔUL36ex2; RCMV2289). Virus stocks were prepared in HFFF-TERTs or HFFF-Tets (for tet-regulated viruses) as described in the Stanton *et al.* study (6). Whole-genome consensus sequences of passage 1 of each RCMV were determined using the Illumina platform as described previously (7).

#### Viral infections

The required volume of viral stock to achieve the multiplicity of infection (MOI) described in the results section was diluted in serum-free DMEM, mixed gently and applied to HFFF-TERTs. Mock infections were performed identically but with DMEM instead of viral stock. Time zero was considered the time at which cells first came into contact with virus. Cells were incubated with virus for 2 h at 37 °C on a rocking platform, and then the medium was replaced with DMEM/FBS.

#### Proteomic Screen

The 48 h degradation screen in this paper was performed in biological duplicate. Cellular lysates from the second 48 h biological replicate were analysed simultaneously with residual lysates from the 12 h degradation screen in the Nightingale *et al.* study (5), thus facilitating a direct comparison.

##### *Infection*

In **Experiment 1**, $3\times10^6$ HFFF-TERTs were plated in DMEM/FBS/PS in 75 $cm^2$ flasks. After 24 h, the medium was replaced by serum-free DMEM containing 4 µg/ml dexamethasone, which has been shown to improve infection efficiency (8, 9). After a further 24 h, the medium was removed and the cells were infected as described above at an MOI of 10. At 36 hours post-infection (hpi), 10 µM MG132 (Merck) or the equivalent volume of DMSO (Sigma-Aldrich) was added to the cells. In **Experiment 2**, $8\times10^5$ HFFF-TERTs were seeded into 25 $cm^2$ flasks, and then treated as in Experiment 1.

##### *Whole Cell Lysate Protein Digestion*

Methods for whole cell lysate protein preparation and digestion, peptide labelling with tandem mass tags, HpRP fractionation, liquid chromatography-mass spectrometry and data analysis are discussed in detail in the Nightingale *et al.* study (5), and are recapitulated below including modifications for the present study.

At 48 hpi, cells were washed twice with PBS, and 500 µl lysis buffer was added (6 M guanidine/50 mM HEPES pH 8.5). Cell lifters (Corning) were used to scrape the cells into lysis buffer, which was then removed to an eppendorf tube, vortexed extensively and sonicated. Cell debris was removed by centrifuging twice at 21,000 g for 10 min.

From this point onward, lysates from the second biological replicate (Experiment 2) were treated identically to residual lysates from a 12 h degradation screen that we performed previously (5). Dithiothreitol (DTT) was added to a final concentration of 5 mM and incubated at room temperature for 20 mins. Cysteine residues were alkylated with 15 mM iodoacetamide and incubated for 20 min at room temperature in the dark. Excess iodoacetamide was quenched with DTT for 15 mins. Samples were diluted with 200 mM HEPES pH 8.5 to 1.5 M guanidine followed by digestion at room temperature for 3 h with LysC protease at a 1:100 protease-to-protein ratio. Samples were further diluted with 200 mM HEPES pH 8.5 to 0.5 M guanidine. Trypsin was then added at a 1:100 protease-to-protein ratio followed by overnight incubation at 37°C. The reaction was quenched with 5% formic acid and centrifuged at 21,000 g for 10 min to remove undigested protein. Peptides were subjected to C18 solid-phase extraction (SPE, Sep-Pak, Waters) and vacuum-centrifuged to near-dryness.

##### *Peptide Labelling with Tandem Mass Tags*

In preparation for TMT labelling, desalted peptides were dissolved in 200 mM HEPES pH 8.5. Peptide concentration was measured by microBCA (Pierce), and 25 µg of peptide was labelled with TMT reagent. TMT reagents (0.8 mg, Thermo Scientific) were dissolved in 43 µl anhydrous acetonitrile and 3 µl were added to each peptide sample at a final acetonitrile concentration of 30% (v/v). Samples were labelled as follows: **Experiment 1:** 48 h mock (TMT 128N), 48 h infection (TMT 129N), 48 h infection / MG132 (TMT 130C). **Experiment 2:** 12 h mock (TMTpro 126), 12 h infection (TMTpro 127C), 12 h infection / MG132 (TMTpro 128N), 48 h mock (TMTpro 131C), 48 h infection (TMTpro 132C), 48 h infection / MG132 (TMTpro 133N). Following incubation at room temperature for 1 h, the reaction was quenched with hydroxylamine to a final concentration of 0.3% (v/v). TMT-labelled samples were combined at a 1:1:1 ratio (**Experiment 1**) or 1:1:1:1:1:1 ratio (**Experiment 2**). The sample was vacuum-centrifuged to near dryness and subjected to C18 SPE (Sep-Pak, Waters). An unfractionated single-shot was analysed initially to ensure similar peptide loading across each TMT channel, thus avoiding the need for excessive electronic normalisation. In **Experiment 1**, data from the single-shot experiment was analysed with data from the corresponding fractions to increase the overall number of peptides quantified. In **Experiment 2**, data from 13 fractions were used in the analysis. Normalisation is discussed in 'Data Analysis' and high pH reversed-phase (HpRP) fractionation is discussed below.

#### *Offline HpRP Fractionation*

TMT-labelled tryptic peptides were subjected to HpRP fractionation using an Ultimate 3000 RSLC UHPLC system (Thermo Fisher Scientific) equipped with a 2.1 mm internal diameter (ID) x 25 cm long, 1.7 µm particle Kinetix Evo C18 column (Phenomenex). Mobile phase consisted of A: 3% acetonitrile (MeCN), B: MeCN, and C: 200 mM ammonium formate pH 10. Isocratic conditions were 90% A/10% C, and C was maintained at 10% throughout the gradient elution. Separations were conducted at 45 °C. Samples were loaded at 200 µl/minute for 5 min. The flow rate was then increased to 400 µl/minute over 5 minutes, after which the gradient elution proceeded as follows: 0-19% B over 10 minutes, 19-34% B over 14.25 minutes, 34-50% B over 8.75 minutes, followed by a 10 min wash at 90% B. UV absorbance was monitored at 280 nm and 15 s fractions were collected into 96 well microplates using the integrated fraction collector. Fractions were recombined orthogonally in a checkerboard fashion, combining alternate wells from each column of the plate into a single fraction, and commencing combination of adjacent fractions in alternating rows. Wells were excluded prior to the start or after the cessation of elution of peptide-rich fractions, as identified from the UV trace. This yielded two sets of 12 combined fractions, A and B, which were dried in a vacuum centrifuge and resuspended in 10 µl MS solvent (4% MeCN/5% formic acid) prior to LC-MS3. 12 set 'A' fractions were used for MS analysis for **Experiment 1** and 12 set 'A' and one set 'B' fractions were used for **Experiment 2**.

## *LC-MS3*

For **Experiment 1,** mass spectrometry data were acquired using an Orbitrap Fusion mass spectrometer (Thermo Fisher Scientific, San Jose, CA). Peptides were separated using a Proxeon EASY-nLC 1000 LC pump equipped with a 75 µm inner diameter microcapillary column packed with 0.5 cm of Magic C4 resin (5 µm, 100 Å, Michrom Bioresources) followed by ~20 cm of GP118 resin (1.8 µm, 120 Å, Sepax Technologies). Peptides were separated using a 3 hr gradient of 6-30% acetonitrile in 0.125% formic acid at a flow rate of 300 nL/min. Each analysis used a MultiNotch MS3-based TMT method (10, 11). The following settings were used: MS1: 400-1400 Th, Quadrupole isolation, 120,000 Resolution, $2\times10^5$ AGC target, 100 ms maximum injection time, ions injected for all parallisable time. MS2: Quadrupole isolation at an isolation width of m/z 0.5, CID fragmentation (NCE 35) with ion trap scanning out in rapid mode from m/z 350, 4000 AGC target, 150 ms maximum injection time, in centroid mode. MS3: In Synchronous Precursor Selection mode, the top 10 MS2 ions were selected for HCD fragmentation (NCE 55) and scanned in the Orbitrap at 60,000 resolution with an AGC target of $5\times10^4$ and a maximum accumulation time of 150 ms. Ions were not accumulated for all parallelisable time. The entire MS/MS/MS cycle had a target time of 3 s. Dynamic exclusion was set to +/- 7 ppm for 90 s. MS2 fragmentation was trigged on precursors $5\times10^3$ counts and above.

For **Experiment 2,** mass spectrometry data were acquired using an Orbitrap Lumos (Thermo Fisher Scientific, San Jose, CA). An Ultimate 3000 RSLC nano UHPLC equipped with a 300 µm ID x 5 mm Acclaim PepMap µ-Precolumn (Thermo Fisher Scientific) and a 75 µm ID x 50 cm 2.1 µm particle Acclaim PepMap RSLC analytical column was used. Loading solvent was 0.1% formic acid (FA), analytical solvent A: 0.1% FA and B: 80% MeCN + 0.1% FA. All separations were carried out at 40°C. Samples were loaded at 5 µl/min for 5 min in loading solvent before beginning the analytical gradient. The following gradient was used: 3-7% B over 3 min, 7-37% B over 173 min, followed by a 4 min wash at 95% B and equilibration at 3% B for 15 min. Each analysis used a MultiNotch MS3-based TMT method (10, 11). The following settings were used: MS1: 380-1500 Th, 120,000 Resolution, $2\times10^5$ automatic gain control (AGC) target, 50 ms maximum injection time. MS2: Quadrupole isolation at an isolation width of m/z 0.7, CID fragmentation (normalised collision energy (NCE) 34) with ion trap scanning in turbo mode from m/z 120, $1.5\times10^4$ AGC target, 120 ms maximum injection time. MS3: In Synchronous Precursor Selection mode the top 10 MS2 ions were selected for HCD fragmentation (NCE 45) and scanned in the Orbitrap at 60,000 resolution with an AGC target of $1\times10^5$ and a maximum accumulation time of 150 ms. Ions were not accumulated for all parallelisable time. The entire MS/MS/MS cycle had a target time of 3 s. Dynamic exclusion was set to ± 10 ppm for 70 s. MS2 fragmentation was trigged on precursors $5\times10^3$ counts and above.

#### Immunoblotting

For the majority of immunoblots in this study, cells were lysed in RIPA buffer (Cell Signalling Technology) containing cOmplete EDTA-free Protease Inhibitor Cocktail (Roche) for 15 minutes at 4°C. For **Figures S2E and S2F**, cells were lysed in 2% SDS in PBS supplemented with 2.5 U/µl benzonase (Sigma) for 30 minutes at 37°C. Supernatants were sonicated and clarified by centrifugation at 14,000 g for 10 minutes. A Bicinchoninic Acid (BCA) assay (Pierce) was used to measure protein concentration according to the manufacturer's instructions. Samples were denatured and reduced with 6× Protein Loading Dye (375 mM Tris pH 6.8, 12% SDS, 30% glycerol, 0.6 M DTT, 0.06% bromophenol blue) for 5 minutes at 95°C. 40 µg (except **Figure 4A**; 50 µg and **Figure S2E**; 20 µg) of protein was separated by SDS-polyacrylamide gel electrophoresis (SDS-PAGE) using Mini-PROTEAN TGX precast gels (Bio-Rad), then transferred to polyvinylidene difluoride (PVDF) membranes (0.45 µm pore) using the Bio-Rad Trans-Blot system. The following primary antibodies were used: anti-MLKL (Cell Signalling Technology, 14993S, 1:1000), anti-IE1/2 (Abcam, ab53495, 1:1000), anti-GAPDH (R&D Systems, MAB5718, 1:10,000), anti-V5 (Invitrogen, R960-25, 1:2000). Secondary antibodies used were IRDye 680RD goat anti-mouse (LI-COR, 925-68070, 1:10,000) and IRDye 800CW goat anti-rabbit (LI-COR, 925-32211, 1:10,000). Fluorescent signals were detected using the Odyssey CLx Imaging System (LI-COR), and images were processed and quantified using Image Studio Lite V5.2 (LI-COR).

#### Plasmid construction

##### *Generation of lentiviral expression vectors*

cDNA was generated from HFFF-TERTs by reverse transcription of RNA using an RNeasy Mini Kit (Qiagen) followed by GoScript Reverse Transcriptase (Promega) according to the manufacturer's protocol. To generate an expression construct for MLKL-HA, primers were designed to recognise the 3' and 5' ends of the MLKL gene and contained flanking Gateway attB sequences to facilitate cloning into pDONR223 using the Gateway system (Thermo Scientific). The reverse primer additionally contained a 6 bp linker region, followed by the coding sequence for an HA tag and a stop codon **(Dataset S3A)**.

For expression of the V5-tagged viral genes, recombinant adenovirus vectors (RAds) were used as a template as described previously (9). Each template expressed a C-terminally V5-tagged gene under the control of the HCMV major immediate early promoter, with a 6 bp linker region between the end of the gene and the tag. To amplify genes from the RAds, primers were designed to recognise the 3' end of the HCMV promoter (forward 'GAW-CMVp-F') and the 3' end of the V5 tag (reverse 'attB2-V5-R') **(Dataset S3A)**. Both primers had flanking Gateway attB sequences.

Spliced UL150A was synthesized as double-stranded DNA fragment (gBlocks®, Integrated DNA Technologies) comprising the viral gene succeeded by a 6 bp linker region, the coding sequence for the V5 tag and then the stop codon, and Gateway attB sequences.

PCR amplification of MLKL-HA and the viral genes was achieved using PfuUltra II fusion HS DNA polymerase (Agilent). PCR products were subsequently purified using a QIAquick Gel Extraction Kit (Qiagen), cloned into the pDONR223 entry vector, and then into the lentiviral destination vector pHAGE-pSFFV (described in the Nightingale *et al.* study (5)) (12). The sequence of MLKL was confirmed as the long, necroptotic isoform and shown to contain two non-disease-causing natural variants (S52T and M169L); this and the confirmed sequences of HCMV genes used in this study are shown in **Dataset S3B**. The UL36 gene sequence contains a single intron. Confirmation of splicing was achieved by reverse transcription of RNA from UL36-expressing

HFFF-TERTs and sequencing of the cDNA **(Dataset S3B)**. The pHAGE-SFFV control vector contains a short randomized DNA sequence (5).

All constructed plasmids were transformed into Alpha-Select Silver Efficiency Competent *E. coli* cells (Bioline) and selected on antibiotic-containing LB agar plates (pDONR223; spectinomycin, pHAGE-pSFFV; ampicillin).

##### *PCR site-directed mutagenesis to generate UL36 point mutants*

In the first round of PCR, two pairs of primers were used in two separate PCRs using strain Merlin UL36-V5 in the pDONR223 vector as a template. In the first reaction, reverse PCR primers encompassing each of the point mutants were combined with the UL36 attB1 fwd primer **(Dataset S3C)**. In the second reaction, forward PCR primers encompassing each of the point mutants were combined with the UL36 V5 attB2 rev primer **(Dataset S3C)**. A second round of PCR joined the two halves of UL36 by using the UL36 attB1 fwd and UL36 V5 attB2 rev primers. Five resulting PCR products encoding each of the different point mutants flanked by attB sites were cloned into pDONR223 and then pHAGE-pSFFV by Gateway cloning, and the sequences were confirmed.

#### Stable cell line production

Lentiviral particles were generated through transfection of HEK293T cells with the lentiviral expression vector and two helper plasmids (VSVg and pCMV.DR8.91, both kindly provided by Professor Paul Lehner), using TransIT-293 (Mirus) according to the manufacturer's instructions. Viral supernatant was harvested 48 h after transfection and diluted to achieve ~30% transduction of target cells, and debris was removed using a 0.22 µM filter (Millipore). 48 h after transduction, cells were subjected to antibiotic selection. Expression of the viral proteins pUL36 and pUL133-pUL150A was confirmed by immunoblotting for the V5 tag or IP-MS (9) (see 'Immunoprecipitation') **(Dataset S3D)**.

#### Immunoprecipitation

Cells were harvested in lysis buffer (50 mM Tris pH 7.5, 300 mM NaCl, 0.5% (v/v) NP40, 1 mM DTT and cOmplete EDTA-free Protease Inhibitor Cocktail (Roche)), tumbled on a rotator for 15 minutes at 4 °C, and then centrifuged at 16,100 g for 15 minutes at 4 °C. Lysates were clarified by filtration through a 0.7 µm filter and incubated for 3 h with immobilised mouse monoclonal anti-V5 or anti-HA agarose resin (Sigma). Samples were washed multiple times with lysis buffer and PBS.

##### *Sample preparation for proteomic analysis*

Confirmation of expression of viral proteins and analysis of SILAC IP data were achieved by using LC-MS/MS. Proteins bound to the resin were eluted twice with 200 µl of 250 µg/ml V5 (Alpha Diagnostic International) or HA (Sigma-Aldrich) peptide in PBS at 37 °C for 30 minutes with agitation. For SILAC-labelled samples, the medium- and heavy-labelled samples were then combined. Proteins were precipitated with 20% Trichloroacetic acid (TCA), washed once with 10% TCA, washed three times with cold acetone and dried to completion. The proteins were then digested in digestion buffer (50 mM Tris-HCl pH 8.5, 10% acetonitrile, 1 mM DTT, 10 µg/ml trypsin) and incubated overnight at 37 °C with agitation. The digestion reaction was quenched with 50% FA, and the peptides were subjected to C18 solid-phase extraction and dried in a centrifugal vacuum. Samples were resuspended in 4% acetonitrile / 5% FA in HPLC grade water and analysed by mass spectrometry on the Orbitrap Lumos as described below.

##### *LC-MS/MS for IP experiments*

Mass spectrometry data were acquired using an Orbitrap Fusion mass spectrometer (Thermo Fisher Scientific, San Jose, CA). Peptides were separated using an Ultimate 3000 RSLC nano UHPLC equipped with a 300 µm ID x 5 mm Acclaim PepMap µ-Precolumn (Thermo Fisher Scientific) and a 75 µm ID x 50 cm 2.1 µm particle Acclaim PepMap RSLC analytical column. Peptides were separated using a 3 hr gradient of 3-37% acetonitrile in 0.125% formic acid at a flow rate of 250 nL/min. The following settings were used: MS1: 350-1500 Th, Quadrupole isolation, 120,000 Resolution, $2\times10^5$ AGC target, 50 ms maximum injection time, ions injected for all parallisable time. MS2: Quadrupole isolation at an isolation width of m/z 0.7, HCD fragmentation (NCE 34) with ion trap scanning out in rapid mode from m/z 120, 10000 AGC target, 250 ms maximum injection time, in centroid mode. Dynamic exclusion was set to +/- 10 ppm for 25 s. MS2 fragmentation was trigged on precursors $5\times10^3$ counts and above.

##### *Sample preparation for immunoblotting*

Proteins bound to the resin were eluted once with 40 µl 2.5 mg/ml V5 or HA peptide at 37 °C for 1 hour with agitation. Eluted proteins were reduced with 6× Protein Loading Dye (Tris 375 mM pH 6.8, 12% SDS, 30% glycerol, 0.6M DTT, 0.06% bromophenol blue) at 95 °C for 5 minutes prior to separation by SDS-PAGE as described above. For the input blot, 2% of the original lysates were removed prior to IP and heated in 6× Loading Dye at 95 °C for 5 minutes prior to SDS-PAGE.

#### Immunofluorescence

1.2×10$^{5}$ HFFF-TERTs were seeded onto 13 mm coverslips (VWR) in a 24-well plate and incubated overnight. Coverslips were washed in the well three times with PBS and then fixed with 4% paraformaldehyde Fixation Buffer (Bio-Rad) for 20 minutes at room temperature. Two more washes with PBS were followed by permeabilisation in 0.1% (v/v) Triton® X-100 (Thermo) / 2% (v/v) FBS in PBS for 4 minutes. Coverslips were then washed twice in each of blocking buffer (10% (v/v) FBS in PBS), PBS and ultrapure water, before blocking for 1 h. The primary antibodies were diluted in blocking buffer and incubated with the coverslips overnight at 4 °C. Coverslips were then washed in blocking buffer, PBS and water as before, and then incubated with secondary antibody in blocking buffer for 1 h. Coverslips were washed again as before, incubated with 1 µg/ml DAPI (Thermo Scientific) in water and washed again before mounting on slides with ProLong™ Gold antifade reagent (Thermo Scientific). The primary antibodies used were anti-HA (Cell Signalling Technology, 3724, 1:50) and anti-V5 (Thermo, MA5-15253, 1:1000). The secondary antibodies used were anti-mouse Alexa Fluor® 488 (Cell Signalling Technology, 4408S, 1:300) and anti-rabbit Alexa Fluor® 647 (Thermo, A31573, 1:300).

#### Cell death assays

All experiments were performed in biological triplicate conducted in parallel and repeated in 2 or 3 completely independent experiments. Input cell numbers and time of stimulation were optimised to ensure that sufficient cells died to facilitate quantitation of statistically significant differences between cell populations, and resulted in absorbance values within the dynamic range of the plate reader. *For assays in the absence of infection,* 96-well plates were seeded with HFFF-TERTs (18,000 cells/well) or immortalised MEFs (6,000 cells/well) then incubated for 24 h. For test cells, the medium was changed to DMEM supplemented with 5% v/v FBS, 30 ng/ml TNFα (R&D systems), 5 µM BV-6 (Selleckchem), or 25 µM z-VAD-fmk (BD Pharmingen) in the presence or absence of 0.5 µM necrosulfonamide (NSA, Merck) or 1.5 µM GSK'872 (Cayman Chemical) for 18 h. Each compound was solubilised in DMSO, and DMSO was added such that its final concentration in each experimental condition was the same. For control cells, the medium was changed to DMEM supplemented with an equivalent concentration of DMSO. 45 mins prior to harvest, half of the control cells were treated with 10 µl 10X Lysis Solution (Promega) to observe lactate dehydrogenase (LDH) release from maximally lysed cells. The other half were used to measure background LDH release from live, untreated cells. Supernatants were harvested and the release of LDH was measured using the CytoTox 96 Non-Radioactive Cytotoxicity Assay (Promega). Average absorbance values derived from culture medium alone were subtracted from each absorbance value from experimental wells. *For assays in the presence of infection*, HFFF-TERTs (14,000 cells/well) were seeded into 96-well plates, incubated for 24 h, and infected with HCMV (MOI = 5). 48 h after infection, the medium was changed as described above to generate populations of cells stimulated for necroptosis, or control cells.

#### Quantification and Statistical Analysis

##### *Data Analysis*

Mass spectra were processed using a Sequest-based software pipeline for quantitative proteomics, ‘‘MassPike’’, through a collaborative arrangement with Professor Steven Gygi’s laboratory at Harvard Medical School. MS spectra were converted to mzXML using an extractor built upon Thermo Fisher’s RAW File Reader library (version 4.0.26). In this extractor, the standard mzxml format has been augmented with additional custom fields that are specific to ion trap and Orbitrap mass spectrometry and essential for TMT quantitation. These additional fields include ion injection times for each scan, Fourier Transform-derived baseline and noise values calculated for every Orbitrap scan, isolation widths for each scan type, scan event numbers and elapsed scan times. This software is a component of the MassPike software platform and is licensed by Harvard Medical School. A combined database was constructed from (a) the human Uniprot database (26th January, 2017), (b) the HCMV strain Merlin Uniprot database, (c) all additional non-canonical HCMV ORFs described by Stern-Ginossar *et al.* (13), (d) a six-frame translation of HCMV strain Merlin filtered to include all potential ORFs of ≥8 amino acids (delimited by stop-stop rather than requiring ATG-stop) and (e) common contaminants such as porcine trypsin and endoproteinase LysC. ORFs from the six-frame translation (6FT-ORFs) were named as follows: 6FT_Frame_ORFnumber_length, where Frame is numbered 1-6, and length is the length in amino acids. The combined database was concatenated with a reverse database composed of all protein sequences in reversed order. Searches were performed using a 20 ppm precursor ion tolerance. Fragment ion tolerance was set to 1.0 Th. TMT tags on lysine residues and peptide N termini (**Experiment 1:** 229.162932 Da; **Experiment 2:** 304.2071) and carbamidomethylation of cysteine residues (57.02146 Da) were set as static modifications, while oxidation of methionine residues (15.99492 Da) was set as a variable modification. Where indicated, data were re-searched using phosphorylation (79.96633 Da) of serine, threonine or tyrosine residues as an additional variable modification. For SILAC analysis, the following variable modifications were used: heavy lysine (8.01420 Da), heavy arginine (10.00827 Da), medium lysine (4.02511 Da), and medium arginine (6.02013 Da). SILAC-only searches were performed in the same manner, omitting the TMT static modification. To control the fraction of erroneous protein identifications, a target-decoy strategy was employed (14). Peptide spectral matches (PSMs) were filtered to an initial peptide-level false discovery rate (FDR) of 1% with subsequent filtering to attain a final protein-level FDR of 1%. PSM filtering was performed using a linear discriminant analysis, as described previously (14). This distinguishes correct from incorrect peptide IDs in a manner analogous to the widely used Percolator algorithm (https://noble.gs.washington.edu/proj/percolator/), though employing a distinct machine-learning algorithm. The following parameters were considered: XCorr, DCn, missed cleavages, peptide length, charge state, and precursor mass accuracy. Protein assembly was guided by principles of parsimony to produce the smallest set of proteins necessary to account for all observed peptides (algorithm described in (14)). Where all PSMs from a given HCMV protein could be explained either by a canonical gene or non-canonical ORF, the canonical gene was picked in preference. In a small number of cases, PSMs assigned to a non-canonical or 6FT-ORF were a mixture of peptides from the canonical protein and the ORF. This most commonly occurred where the ORF was a 5'-terminal extension of the canonical protein (thus meaning that the smallest set of proteins necessary to account for all observed peptides included the ORFs alone). In these cases, the peptides corresponding to the canonical protein were separated from those unique to the ORF, generating two separate entries. In a single case, PSM were assigned to the 6FT-ORF 6FT_6_ORF1202_676aa, which is a 5'-terminal extension of the non-canonical ORF ORFL147C. The principles described above were used to separate these two ORFs. Proteins were quantified by summing TMT reporter ion counts across all matching peptide-spectral matches using ''MassPike'', as described previously (10). Briefly, a 0.003 Th window around the theoretical m/z of each reporter ion was scanned for ions and the maximum intensity nearest to the theoretical m/z was used. The primary determinant of quantitation quality is the number of TMT reporter ions detected in each MS3 spectrum, which is directly proportional to the signal-to-noise (S:N) ratio observed for each ion. Conservatively, every individual peptide used for quantitation was required to contribute sufficient TMT reporter ions so that each on its own could be expected to provide a representative picture of relative protein abundance (10). An isolation specificity filter with a cut-off of 50% was additionally employed to minimise peptide co-isolation (10). Peptide-spectral matches with poor quality MS3 spectra (a combined S:N ratio of less than 120 (**Experiment 1**) or 240 (**Experiment 2**) across all TMT reporter ions) or no MS3 spectra at all were excluded from quantitation. Peptides meeting the stated criteria for reliable quantitation were then summed by parent protein, in effect weighting the contributions of individual peptides to the total protein signal based on their individual TMT reporter ion yields. Protein quantitation values were exported for further analysis in Excel. For protein quantitation, reverse and contaminant proteins were removed, then each reporter ion channel was summed across all quantified proteins and normalised assuming equal protein loading across all channels. To combine data from 48 h experiments 1 and 2, fractional TMT signals compared to the sum across all three assessed conditions were used. For example, for protein PCDHGB5, three fractions were calculated for each experiment: mock/(sum(mock,HCMV,HCMV+MG)), HCMV/(sum(mock,HCMV,HCMV+MG)), HCMV+MG/(sum(mock,HCMV,HCMV+MG)). Replicate fractions from each experiment were averaged for display in figures, and for the purposes of fold change calculation. This effectively corrected for differences in the numbers of peptides observed per protein. For the TMT-based experiments, both normalised S:N values and fractional TMT signals are presented in **Dataset S1** ('Data' worksheet). For proteins only quantified in one replicate, the '48 h average' plot in this dataset shows the data from the replicate in which the protein was quantified; otherwise, the average and range of the two fractional TMT signals. Data are similarly displayed in **Figures 1D-E**. Although peptides were assigned appropriately to HLA-A alleles, it was not possible to assign peptides confidently to only two HLA-B or HLA-C alleles, and signal:noise values were further summed for each of these alleles to give a single combined result for HLA-B or HLA-C.

**Figure 1 and S1B**. K-means clustering was performed using Cluster 3.0 (Stanford University) and visualised using Java Treeview (http://jtreeview.sourceforge.net).

**Figure S5**. Alignment of pUL36 amino acid sequences with performed using Clustal Omega (15) (https://www.ebi.ac.uk/Tools/msa/clustalo/) by EMBL-EBI.

##### *Statistical analysis*

**Figures 1, S1A and S4A**. The method of Significance B was used to estimate the p-value that the fold downregulation or rescue was significantly different to 1 (16). Values were calculated and corrected for multiple hypothesis testing using the method of Benjamini-Hochberg in Perseus version 1.5.2.20 (16).

**Figure 2**. P-values for enrichment in immunoprecipitations were estimated using the method of Significance A and corrected for multiple hypothesis testing in Perseus version 1.5.2.20 (16).

**Figures 3 and 4**. P-values were calculated using a Student's two-sample, two-tailed t-test in R version 3.4.1 (17). A Shapiro-Wilk test was used to determine that the data was normally distributed and therefore valid for analysis by a t-test. Homogeneity of variance was confirmed using a Bartlett's test.

#### Data and materials availability statement

The mass spectrometry proteomics data have been deposited to the ProteomeXchange Consortium (http://www.proteomexchange.org/) via the PRIDE partner repository (18) with the dataset identifier PXD017279. All materials described in this manuscript, and any further details of protocols employed, can be obtained on request from the corresponding author by email to.

**Supplementary Information Figures**

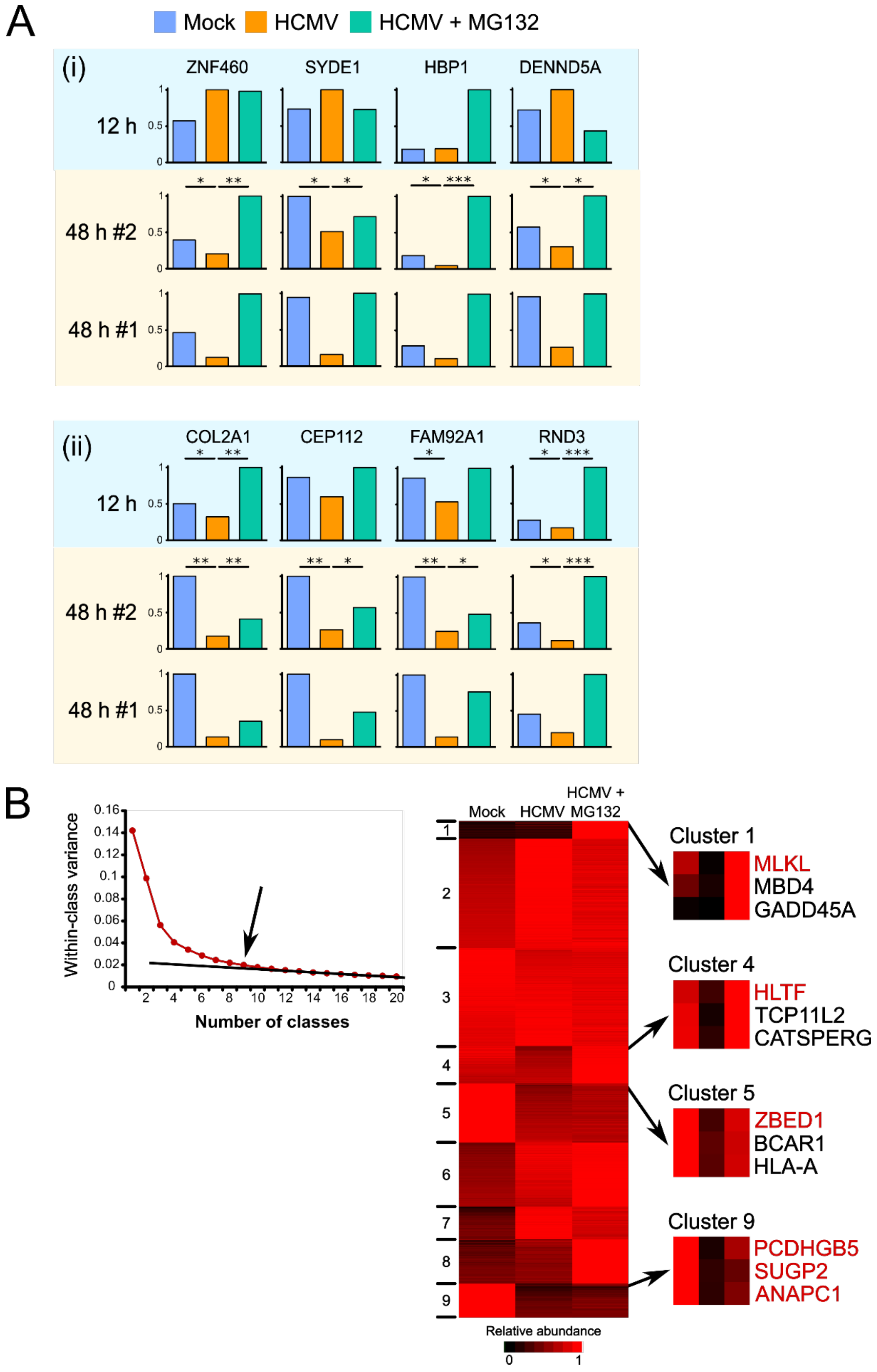

**Figure S1. (A)** Comparison of 12 and 48 h degradation data for individual proteins. A direct contrast can be made between 12 h data and the second 48 h biological replicate, as both samples were simultaneously digested into peptides, labelled with TMT reagents and analysed. Data for the first 48 h replicate are also shown for comparison. 12 h data were derived by re-analysis of residual protein samples from the Nightingale *et al.* study (5). (i) Four proteins degraded at 48 h but not at 12 h. (ii) Four proteins degraded at 48 h, but insufficiently degraded at 12 h to pass filtering criteria in the Nightingale *et al.* study (5). P-values are shown above the data for 48 h experiment 2 and were derived from ratios generated from averaged data from experiments 1 and 2 using Significance B values **(Figure 1B)** $*p<0.05$, $**p<0.001$, $***p<1\times10^{-7}$. A full list of proteins degraded late in infection can be found in **Dataset S1**, including details of which are also degraded early. **(B)** K-means-based hierarchical cluster analysis of all human proteins quantified **(Figure 1B)**. K-means clustering with 1-20 classes was used to assess the summed distance of each protein from its cluster centroid. While this summed distance necessarily became smaller as more clusters were added, the rate of decline decreased with each added group, eventually settling at a fairly constant rate of decline that reflected over-fitting; clusters added prior to this point reflected underlying structure in the protein data, while clusters subsequently added through over-fitting were not informative. The point of inflexion fell at or after nine classes, indicating that there were at least nine classes of proteins displaying different expression profiles across the protein samples. The panels on the right show examples of proteins belonging to classes 1, 4, 5 and 9, which contain the 7 proteins degraded early and late during infection **(Figure 1C)**. A full list of proteins in each cluster can be found in **Dataset S1C**.

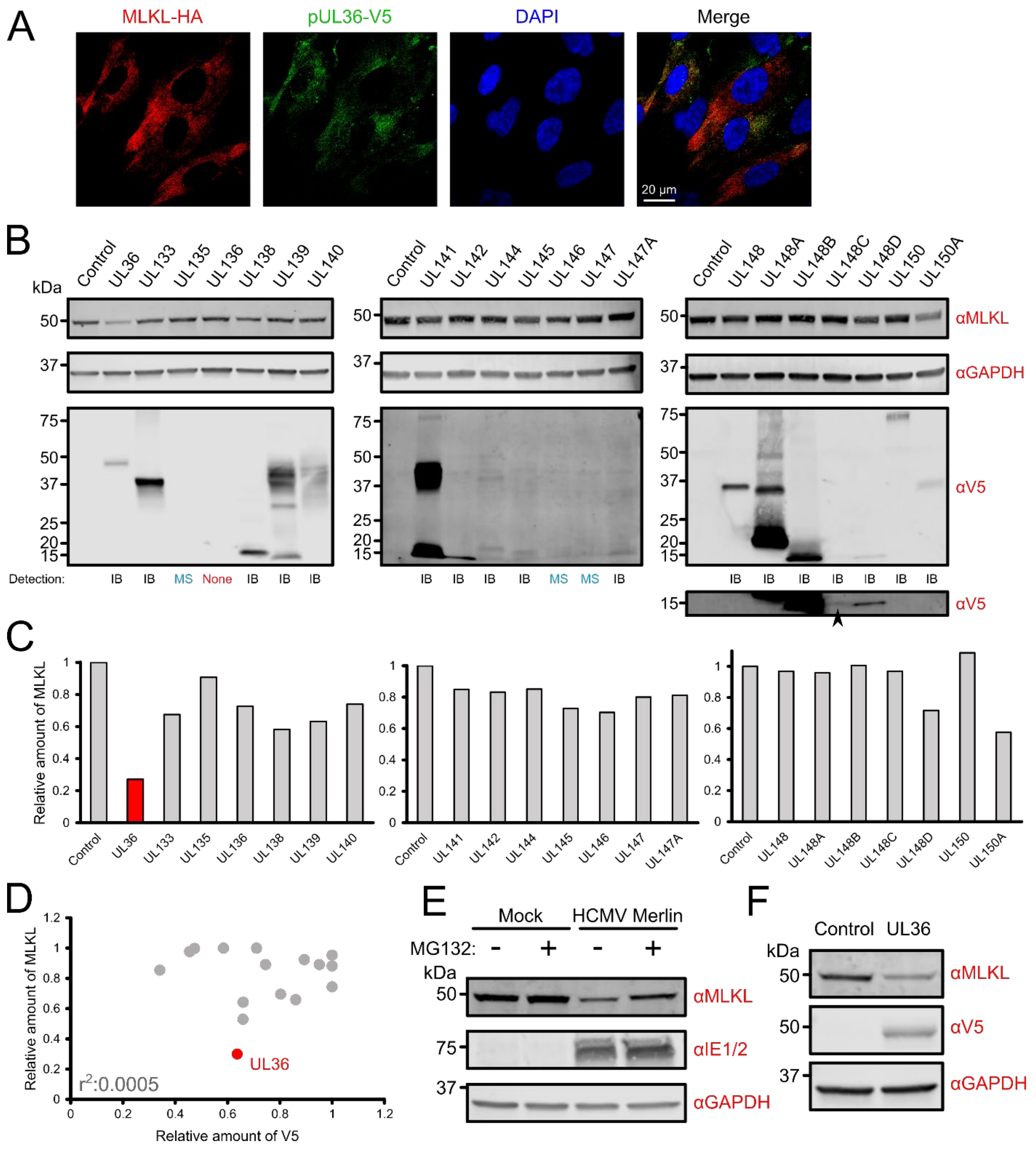
A
MLKL-HA
pUL36-V5
DAPI
Merge
20 µm
B
kDa
Control
UL36
UL133
UL135
UL136
UL138
UL139
UL140
UL141
UL142
UL144
UL145
UL146
UL147
UL147A
UL148
UL148A
UL148B
UL148C
UL148D
UL150
UL150A
αMLKL
αGAPDH
αV5
Detection:
IB
MS
None
C
Relative amount of MLKL
D
r²:0.0005
UL36
Relative amount of V5
E
Mock
HCMV Merlin
MG132:
αIE1/2
F
Control
UL36

**Figure S2. (A)** Immunofluorescence demonstrated diffuse cytoplasmic colocalisation between MLKL-HA and pUL36-V5 in stably co-transduced HFFF-TERTs. **(B)** Immunoblot showing that pUL36 is sufficient to downregulate MLKL. Expression of 17/21 V5-tagged viral proteins was confirmed by anti-V5 immunoblot. Expression of pUL135, pUL146 and pUL147 was confirmed by mass spectrometry (9). pUL136 was not detected by either method; further confirmation that pUL136 does not affect MLKL levels is required, although this protein was not detected in the SILAC IP of MLKL-HA in HCMV-infected cells (**Figure 2C**). A faint band corresponding to pUL148C could be detected upon over-exposure of the blot (bottom right panel). **(C)** Densitometry analysis of MLKL expression normalised by GAPDH expression in each cell line. Expression of UL36 was sufficient to reduce the level of MLKL by 3.7-fold. MLKL expression was not modulated more than 2-fold by other proteins in the UL/b' region. **(D)** Despite substantial variation in the level of expression of some of the viral proteins, this did not correlate with relative MLKL abundance. For every cell line in which V5 was detectable by immunoblotting, the relative abundances of MLKL (adjusted to GAPDH) and the V5-tagged viral protein (adjusted to GAPDH) were calculated. To prevent confounding effects from systematic expression level differences between each of the three blots, values derived from each blot were normalised to the maximum level of MLKL and V5 expression prior to comparison. No correlation between MLKL and V5 expression was observed. **(E)** Immunoblot confirming that MLKL is degraded by HCMV strain Merlin in cells solubilised in 2% SDS as well as in cells lysed in RIPA buffer **(Figure 2B)**. The level of MLKL was partially recovered by addition of MG132, indicating active degradation (MOI = 5, 48 h infection). MG132 at a final concentration of 10 μM or an equivalent volume of DMSO was added to media for the last 12 h of infection. **(F)** Immunoblot confirming that MLKL is downregulated by pUL36 in cells solubilised in 2% SDS as well as in cells lysed in RIPA buffer **(Figure 2E)**.

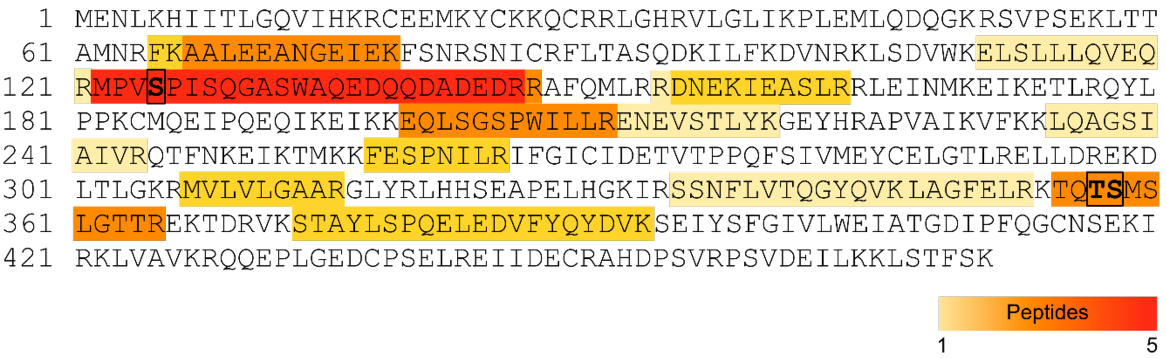


**Figure S3.** pUL36 interacts with unphosphorylated MLKL. Amino acid sequence of MLKL, with the peptides identified in the pUL36-V5 IP **(Figure 2D)** highlighted in different colours according to the frequency of identification. Thr357 and Ser358 are phosphorylated by RIP3 during TNFα-stimulated necroptosis (19). Ser125 is phosphorylated in a cell-cycle dependent manner but its effect on MLKL function is unknown (20, 21). No phospho-MLKL peptides were identified.

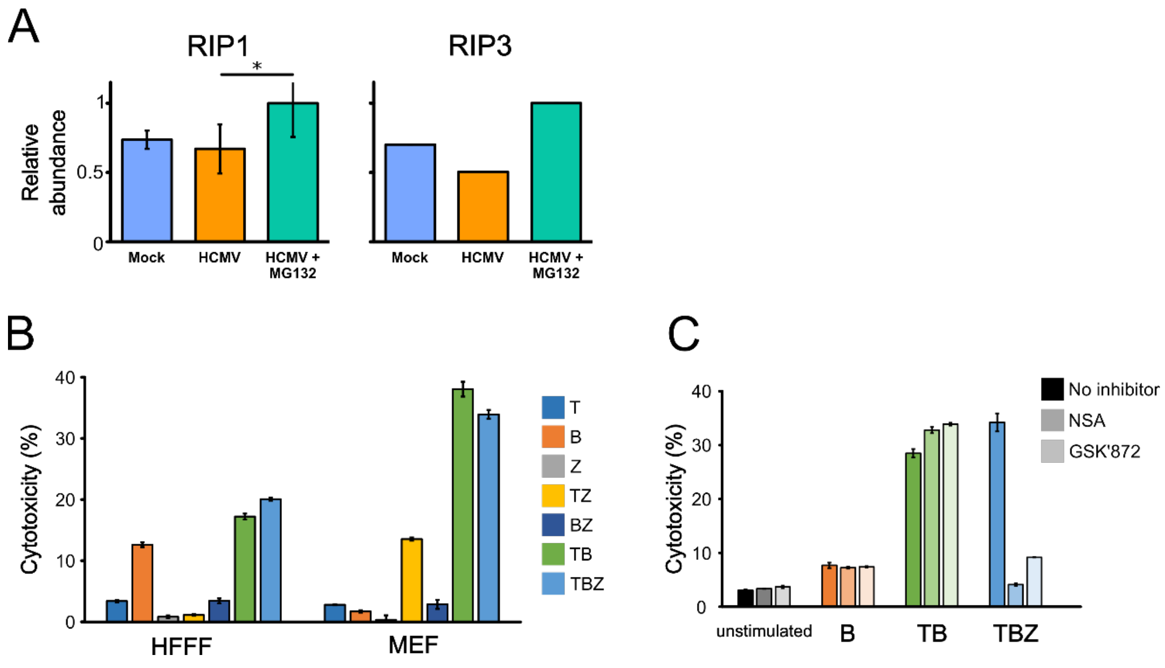


**Figure S4**. **(A)** Expression of RIP1 and RIP3 in HFFF-TERTs quantified by proteomics (the experimental setup is described in **Figure 1**). RIP3 was only detected in the first of two biological replicates. *$p<0.05$. **(B)** Cell death in HFFF-TERTs is stimulated by BV6 (B) alone, TNFα + BV6 (TB), and TNFα + BV6 + Z-VAD-fmk (TBZ). In addition to apoptosis stimulated by TB and necroptosis stimulated by TBZ, HFFFs also underwent cell death in response to B alone, which has been described in other cell types (22). Cell death in MEFs was additionally stimulated by TZ, which has been observed in other mouse fibroblast cell lines (23). Each treatment was applied for 18 h. Error bars show standard error of the mean (SEM). Data are representative of three independent experiments. **(C)** Cell death in HFFF-TERTs stimulated by TBZ is RIP3- and MLKL-dependent, whereas B or TB stimulate pathways independent of RIP3 and MLKL. Cells were treated for 18 h in the presence or absence of necrosulfonamide (NSA) or GSK’872. Application of NSA and GSK’872 in the absence of T, B or Z indicated that the inhibitors did not impact cell viability.

```
 AD169  MDDLRDTLMAYGCIAIRAGDFNGLNDFLEQECGTRLHVAWPERCFIQLRSRSALGPFVGK 60
Merlin  MDDLRDTLMAYGCIAIRAGDFNGLNDFLEQECGTRLHVAWPERCFIQLRSRSALGPFVGK 60
        ************************************************************

 AD169  MGTVCSQGAYVCCQEYLHPFGFVEGPGFMRYQLIVLIGQRGGIYCYDDLRDCVYELAPTM 120
Merlin  MGTVCSQGAYVCCQEYLHPFGFVEGPGFMRYQLIVLIGQRGGIYCYDDLRDCIYELAPTM 120
        ****************************************************:*******

 AD169  KDFLRNGFRHRDHFHTMRDYQRPMVQYDDYWNAVMLYRGDVESLSAEVTKRGYASYTIDD 180
Merlin  KDFLRHGFRHCDHFHTMRDYQRPMVQYDDYWNAVMLYRGDVESLSAEVTKRGYASYSIDD 180
        *****:**** *********************************************:***

 AD169  PFDECPDTHFAFWTHNTEVMKFKETSFSVVRAGGSIQTMELMIRTVPRITCYHQLLGALG 240
Merlin  PFDECPDTHFAFWTHNTEVMKFKETSFSVVRAGGSIQTMELMIRTVPRITCYHQLLGALG 240
        ************************************************************

 AD169  HEVPERKEFLVRQYVLVDTFGVVYGYDPAMDAVYRLAEDVVMFTCVMGKKGHRNHRFSGR 300
Merlin  HEVPERKEFLVRQYVLVDTFGVVYGYDPAMDAVYRLAEDVVMFTCVMGKKGHRNHRFSGR 300
        ************************************************************

 AD169  REAIVRLEKTPTCQHPKKTPDPMIMFDEDDDDELSLPRNVMTHEEAESRLYDAITENLMH 360
Merlin  REAIVRLEKTPTCQHPKKTPDPMIMFDEDDDDELSLPRNVMTHEEAESRLYDAITENLMH 360
        ************************************************************

 AD169  CVKLVTTDSPLATHLWPQELQALCDSPALSLCTDDVEGVRQKLRARTGSLHHFELSYRFH 420
Merlin  CVKLVTTDSPLATHLWPQELQALCDSPALSLCTDDVEGVRQKLRARTGSLHHFELSYRFH 420
        ************************************************************

 AD169  DEDPETYMGFLWDIPSCDRCVRRRRFKVCDVGRRHIIPGAANGMPPLTPPHVYMNN     476
Merlin  DEDPETYMGFLWDIPSCDRCVRRRRFKVCDVGRRHIIPGAANGMPPLTPPHAYMNN
        ***************************************************.****
```

**Figure S5.** Amino acid sequence alignment of pUL36 from HCMV strains Merlin and AD169 using Clustal Omega by EMBL-EBI (15). The five amino acid sequence differences are highlighted. '*' indicates positions which have an identical, fully conserved residue. ':' indicates a substitution with a residue with strongly similar properties. '.' indicates a substitution with a residue with weakly similar properties.

**Supplementary Information Legends for Datasets**

**Dataset S1 (separate .xls file).** Interactive spreadsheet of all TMT-based proteomic data in this paper. The 'Plotter' worksheet generates graphs of protein abundance for any human or viral protein quantified across each experiment. The 'Data' worksheet shows minimally annotated protein data, for which the only modifications are formatting, deletion of contaminants, normalisation and reassignment of non-canonical HCMV ORFs. The 'K-means clusters' worksheet shows the human proteins belonging to each cluster from **Figure S1B**.

**Dataset S2 (separate .xls file).** Data from the MLKL-HA (denoted 'HA') and pUL36-V5 (denoted 'V5') SILAC immunoprecipitations.

**Dataset S3 (separate .xls file). (A)** Oligonucleotides employed in PCR amplification of MLKL-HA and the V5-tagged viral genes used in the study. **(B)** Confirmed sequences of MLKL-HA and the viral genes UL36 and UL133-150A. **(C)** Oligonucleotides employed for site-directed mutagenesis of UL36. **(D)** Method of cloning and verification of expression of each of the viral genes UL36 and UL133-150A.

**Supplementary Information References**

86(14):7150–7158.
11. McAlister GC, et al. (2012) Increasing the multiplexing capacity of TMT using reporter ion isotopologues with isobaric masses. *Anal Chem* 84:7469–7478.
12. Hartley JL, Temple GF, Brasch MA (2000) DNA cloning using in vitro site-specific recombination. *Genome Res* 10:1788–1795.
13. Stern-Ginossar N, et al. (2012) Decoding human cytomegalovirus. *Science (80- )* 338(6110).
14. Huttlin EL, et al. (2010) A tissue-specific atlas of mouse protein phosphorylation and expression. *Cell* 143(7):1174–1189.
15. Sievers F, et al. (2011) Fast, scalable generation of high-quality protein multiple sequence alignments using Clustal Omega. *Mol Syst Biol* 7(1):539.
16. Cox J, Mann M (2008) MaxQuant enables high peptide identification rates, individualized p.p.b.-range mass accuracies and proteome-wide protein quantification. *Nat Biotechnol* 26(12):1367–1372.
17. R Core Team (2018) R: A language and environment for statistical computing. Available at: https://www.r-project.org/.
18. Vizcaíno JA, et al. (2016) 2016 update of the PRIDE database and its related tools. *Nucleic Acids Res* 44(D1):D447–D456.
19. Wang H, et al. (2014) Mixed lineage kinase domain-like protein MLKL causes necrotic membrane disruption upon phosphorylation by RIP3. *Mol Cell* 54(1):133–146.
20. Dephoure N, et al. (2008) A quantitative atlas of mitotic phosphorylation. *PNAS* 105(31):10762–10767.
21. Daub H, et al. (2008) Kinase-selective enrichment enables quantitative phosphoproteomics of the kinome across the cell cycle. *Mol Cell* 31(3):438–448.
22. Christofferson DE, Li Y, Yuan J (2014) Control of life-or-death decisions by RIP1 kinase. *Annu Rev Physiol* 76(1):129–150.
23. Ros U, et al. (2017) Necroptosis execution is mediated by plasma membrane nanopores independent of calcium. *Cell Rep* 19(1):175–187.
